# Dynamic remodeling of centrioles and the microtubule cytoskeleton in the lifecycle of chytrid fungi

**DOI:** 10.1101/2025.01.03.631223

**Authors:** Alexandra F. Long, Krishnakumar Vasudevan, Andrew J.M. Swafford, Claire M. Venard, Jason E. Stajich, Lillian K. Fritz-Laylin, Jessica L. Feldman, Tim Stearns

**Affiliations:** Department of Biology, Stanford University, Stanford, CA, USA; Department of Biology, University of Kentucky, Lexington, KY, USA; Department of Biology, Middlebury College, Middlebury, VT, USA; Department of Microbiology and Plant Pathology, University of California Riverside, Riverside, CA, USA; Department of Biology, University of Massachusetts Amherst, Amherst, MA, USA; Laboratory of Cellular Dynamics, Rockefeller University, New York, NY, USA

## Abstract

Cells reorganize in space and time to move and divide – complex behaviors driven by their internal cytoskeleton. While we have substantial knowledge of the molecular parts and rules of cytoskeletal assembly, we know less about how structures are remodeled, for example to interconvert centrioles from the ciliary base to the centrosome. To study this in an evolutionary context we use the chytrid fungus, *Rhizoclosmatium globosum*, a member of the zoosporic fungi which have centrioles and cilia, lost in other fungal lineages. Chytrids undergo reorganization of their microtubule cytoskeleton as they cycle from zoospore to multinucleated coenocyte. We use comparative bioinformatics, RNA sequencing, and expansion microscopy to map the microtubule cytoskeleton over the chytrid lifecycle. We find that when zoospores encyst, cilia are retracted into the cytoplasm and degraded, and centrioles detach but are protected from degradation. A shortened proximal centriole then forms the mitotic centrosome and ultimately elongates to form cilia at the end of the mitotic cycles, driven by a conserved transcriptional program. Thus, structural remodeling of the chytrid centriole is coupled temporally to ciliated stages rather than mitotic cycles, which may serve as a mechanism to tune microtubule organization to meet the needs of different lifecycle stages.

## Introduction

Cells organize and rearrange their contents in space and time to perform complex behaviors such as dividing and swimming, driven by their dynamic cytoskeleton. The microtubule cytoskeleton is patterned in the cell by microtubule organizing centers (MTOCs). Defects in the spatial and temporal patterning of the microtubule cytoskeleton can compromise cell division, motility, and signaling, causing diseases from ciliopathies to cancer (Reiter and Leroux, 2017).

One important and highly conserved microtubule structure in the eukaryotic cell is the cilium/flagellum, a long hair-like protrusion that has roles in signaling and motility. The base of the cilium is a barrel-shaped centriole made from microtubules, from which emerges the axoneme, a long and highly patterned bundle of specialized microtubules. Organisms or cell types with cilia often alternate states for the centriole between serving as the base of the cilium and as the center of the centrosome. Centrosomes are microtubule organizing centers which have a pair of centrioles surrounded by a matrix of pericentriolar material (PCM) that nucleates and organizes arrays of microtubules in interphase and the spindle during mitosis. For centrioles to play these dual roles in the cell as part of cilia and centrosomes, they must be uncoupled and remodeled over time (Kalbfuss and Gönczy, 2023). While decades of work have focused on the parts list and rules for building cytoskeletal structures, we know comparatively less about the molecular and structural basis for how MTOCs are remodeled to enable distinct cell states and functions. In part this is due to the fact that much of what we know about MTOC remodeling comes from the study of animal centrosomes in cycling cells *in vitro*. To understand more about how cells remodel MTOCs *in vivo*, we need organisms and cell types that readily undergo cilium and centrosome remodeling events during their cell cycle or lifecycle.

Zoosporic fungi are a sister group to the Dikarya (mushrooms, molds and yeasts) and retain motile cilia and centrioles that were lost in other fungal lineages (Powell, 1980, 2017). Thus chytrids, as zoosporic fungi, occupy a key phylogenetic position to shed light onto MTOC biology and evolution (Yubuki and Leander, 2013; Ito and Bettencourt-Dias, 2018). These unicellular organisms have a complex and highly conserved microtubule cytoskeleton that they extensively remodel over a short synchronous lifecycle as they develop from a motile zoospore to a stationary multinucleated sporangium. They reel their long cilium into their cell body then degrade this massive structure by an unknown mechanism (Koch, 1968; Laundon *et al*., 2022). They then remodel their centrioles to form a mitotic MTOC at the spindle poles and go through multiple mitoses without cell division to create a large multinucleated coenocytic sporangium. In this shared cytoplasm they build one new cilium per nucleus before subdividing into new zoospores that are released to the environment.

Historically chytrid cell biology has focused extensively on taxonomy, and only recently have there been efforts to map molecular changes over the lifecycle (Rosenblum *et al*., 2008). Importantly, chytrids play critical ecological roles in aquatic ecosystems (Gleason Frank H. *et al*., 2017) and parasitize organisms from amphibians to algae (Longcore *et al*., 1999; Ito and Bettencourt-Dias, 2018; Venard *et al*., 2020) and this parasitism is intimately related to the remodeling of the chytrid cytoskeleton. Tools for genetic transformation are just starting to be developed for a few chytrid species (Medina *et al*., 2020; Swafford *et al*., 2020; Kalinka *et al*., 2023) and it has been challenging to visualize chytrid cells with immunofluorescence due to the small size of zoospores and the complex fungal cell wall of the sporangia. Thus, we currently lack molecular and structural understanding of how chytrids execute these dramatic cytoskeletal changes in normal lifecycles and in pathogenic contexts. How is the cilium-centriole complex disassembled during ciliary retraction and how is it reformed during ciliogenesis in the sporangium?

To address these questions, we use the non-pathogenic chytrid fungus *Rhizoclosmatium globosum* (*Rg*) together with RNA sequencing and high-resolution ultrastructure expansion microscopy to characterize microtubule reorganization during the coenocytic life cycle. Many chytrid species retain a complex actin cytoskeleton that enables amoeboid crawling in addition to swimming with a motile cilium (Prostak *et al*., 2021). *Rg* has a reduced actin cytoskeleton and does not crawl, has a highly synchronous and rapid lifecycle, making it an excellent model (Laundon and Cunliffe, 2021; Laundon *et al*., 2022) for specifically studying remodeling of the microtubule cytoskeleton. Here we find that chytrids have complex microtubule organizing centers at different lifecycle stages and dynamically remodel centrioles and MTOCs as they transit between ciliogenesis, interphase, and mitosis.

## Results and Discussion

### Developing tools to study chytrid fungi

To develop chytrid fungi as a system for studying centriole and microtubule organizing center remodeling, we first characterized their lifecycle. We chose the chytrid species *Rhizoclosmatium globosum* because it has a sequenced genome, a synchronous 20 hour lifecycle, and exhibits representative chytrid cell architecture and development (Berger *et al*., 2005; Mondo *et al*., 2017; Powell, 2017) (Figure 1A,B). Using *Rg* strain JEL800 we observed the progression of the life cycle, from initial zoospore formation to the release of new zoospores, using light microscopy. A cartoon schematic (Figure 1A) illustrates the life cycle, starting from the motile zoospore, which undergoes cilium retraction and centriole detachment, followed by mitotic divisions without cytokinesis, forming a multinucleate sporangium. Hyphal-like rhizoids anchor the sporangium to the substrate as it enlarges and develops. Eventually after building new cilia from each pair of centrioles, the sporangium cellularizes into separate zoospores that are then released. Phase contrast microscopy (Figure 1B) confirmed the temporal dynamics and synchrony of this process in this strain, with zoospore plating leading to sporangium maturation and a gradual increase in sporangium diameter followed by zoospore release.

**Figure 1.**
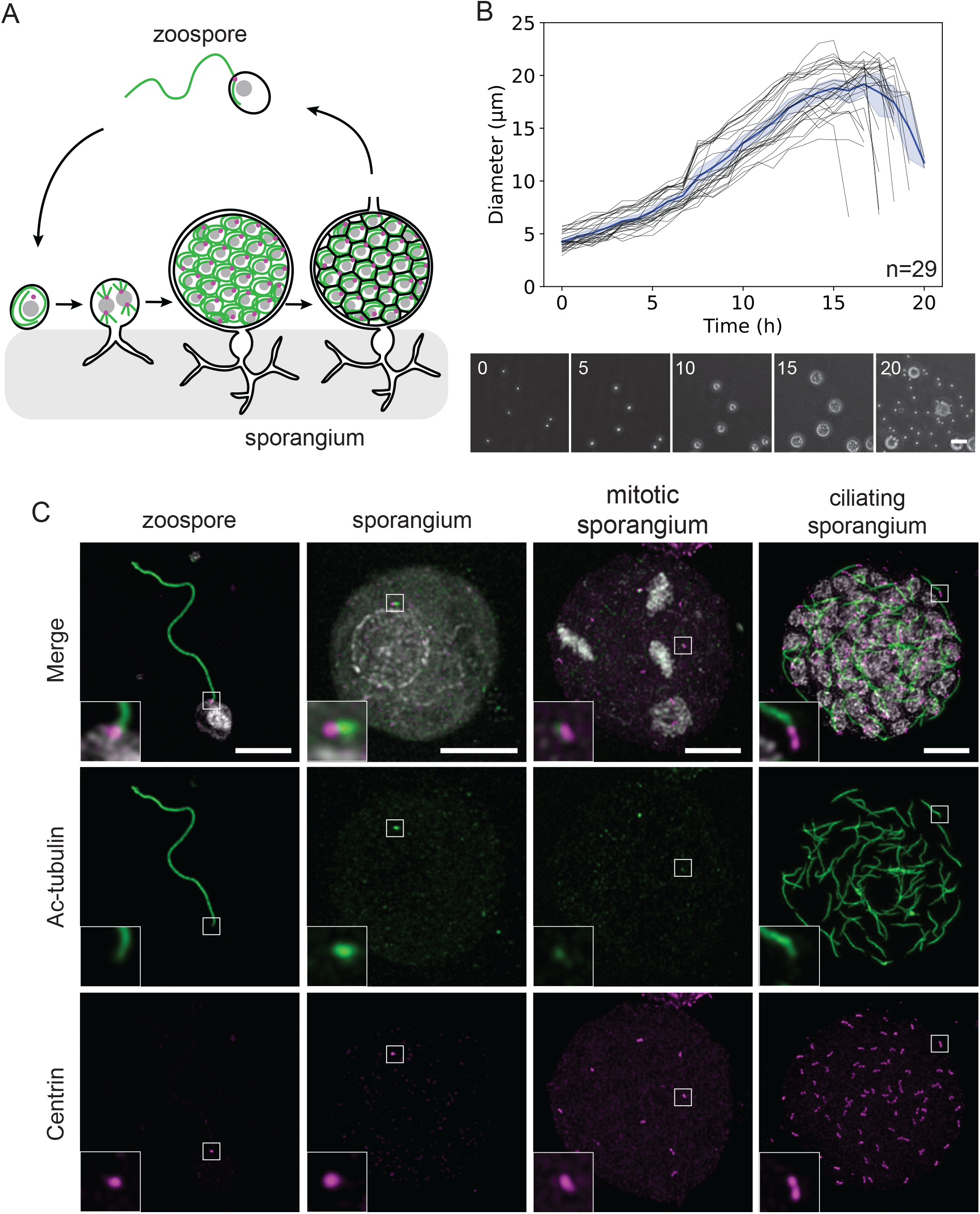
*R.hizoclosmatium globosum* has a synchronous coenocytic lifecycle. A) Cartoon of the lifecycle of a chytrid fungus beginning from a motile zoospore (top) that retracts its cilium (green) into the cell body and detaches centrioles (pink) before progressing through mitotic cycles without cytokinesis. Rhizoids at the base of the sporangium attach to the substrate. Ultimately the coenocytic sporangium contains dozens of nuclei and centrioles that undergo ciliogenesis in the shared cytoplasm before cellularizing into individual zoospores. B) Timecourse of *R. globosum* lifecycle imaged with phase contrast microscopy from zoospore plating to new zoospore release. Scale bar 100 μm. Quantification of raw (black) and average (blue) sporangium diameter over the lifecycle showing synchronous development (n=29 sporangia from 3 independent experiments). C) Immunofluorescence characterization of subcellular structures over the chytrid lifecycle (DAPI; white, acetylated-tubulin; green, centrin; magenta). The zoospore has a cilium and centriole while the sporangium has degraded the ciliary axoneme but not the centriole and increases in size as it matures and enters mitosis. This example shows synchronous metaphase spindles of the second mitosis. Synchronous ciliogenesis occurs after the end of the mitotic cycles. Scale bar, 10 μm (insets 2x enlarged).

Immunofluorescence is an essential tool for studying the cytoskeleton, but it has been technically challenging in chytrids due to the chitin-based cell wall present during much of the life cycle. Thus, we developed an optimized protocol for immunofluorescence using chitinase treatment and physical freeze-cracking of *Rg* sporangia. This protocol enables labeling of the *Rg* microtubule cytoskeleton at all life cycle stages using immunofluorescence rather than electron microscopy (Figure 1C). We characterized markers for centrioles and cilia in chytrid zoospores and sporangia. First we examined a cilium and centriole marker using an antibody against acetylated-tubulin, a post-translational modification of tubulin that decorates the stable microtubules populations (Piperno *et al*., 1987). As is seen in other systems, in *Rg* axonemal microtubules are highly acetylated (Figure 1C) in comparison to other microtubules and acetylation is higher on the centrioles at the poles of the mitotic spindle, rather than on the microtubules of the spindle itself in mitotic sporangia. To specifically localize centrioles at each stage, we used an antibody against centrin, a well-conserved protein that plays a role in centriole duplication and centriole tethering in diverse eukaryotes (Taillon *et al*., 1992; Levy *et al*., 1996; Zhang and He, 2012). As predicted from its genome, *Rg* contains an ortholog of centrin that localizes (Figure 1C) at and surrounding centrioles in both zoospores and sporangia. Thus chytrids have highly conserved components of the microtubule cytoskeleton and we are able to identify clear molecular markers to distinguish the microtubules of the centriole and cilium at different stages of the chytrid lifecycle.

### Bioinformatic characterization of chytrid centriole associated genes

Here we expand what is known of the conservation of core motile cilium components in chytrids to include components of overall microtubule organization (Hodges *et al*., 2010; Ito and Bettencourt-Dias, 2018; Galindo *et al*., 2021). In many cells, centrioles transit between serving as the center of the centrosome and the base of the cilium. Remodeling the centriole to form these different MTOCs requires distinct regulators and functionalization. In most fungal lineages, centrioles and cilia have been lost and cells instead have a non-centrosomal MTOC called the spindle pole body which has few components in common with animal centrosomes (Ito and Bettencourt-Dias, 2018). Given that the chytrid lineages of fungi have retained centrioles and cilia, we determined which microtubule organizing components they possessed. To address this, we took a comparative approach using OrthoFinder (Emms and Kelly, 2015, 2019) and a set of well-established genes associated with mammalian centrioles and PCM as well as fungal spindle pole bodies across all fungal lineages that have a zoospore with motile cilia (Figure 2). We find that zoosporic fungi have many conserved genes with metazoans involved in core centriole structure as well as MTOC and basal body functionalization.

**Figure 2.**
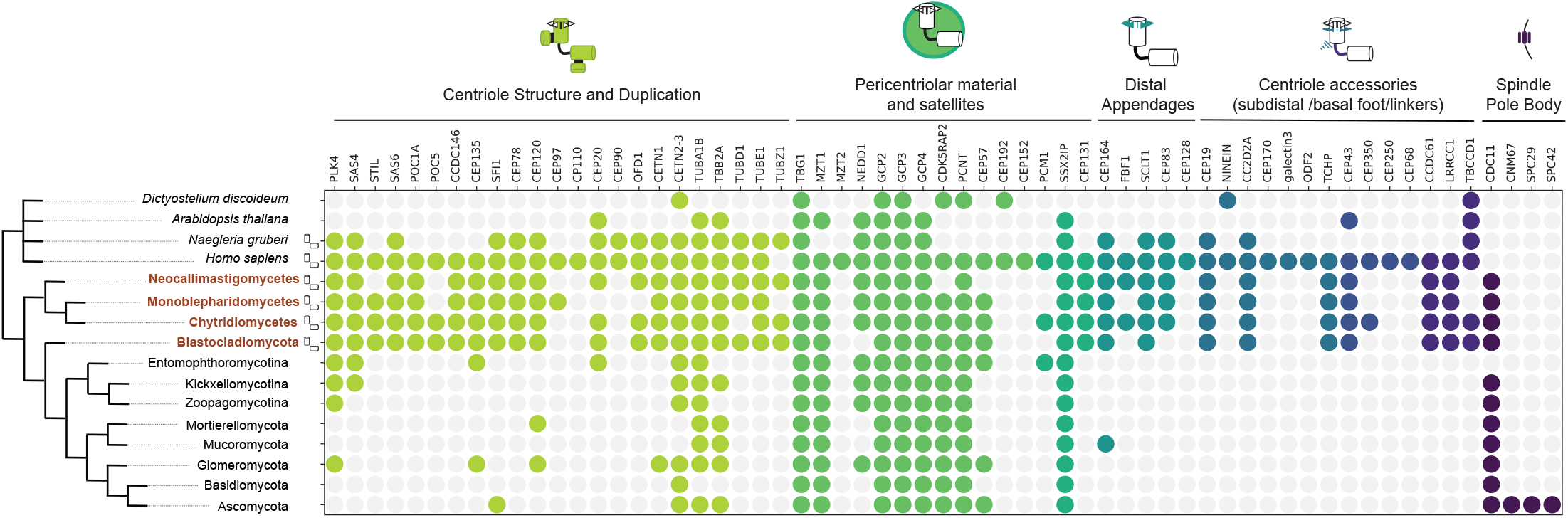
Broad conversation of centriole and centrosome associated genes in zoosporic fungi. Orthologs of genes associated with different centriole and centrosome features across (left) four outgroups (gray, top) and fungal phyla and subphyla (bottom). Each colored circle denotes that there was at least one ortholog assignment in species of the associated fungal clade compared to four outgroups that include two species with (*H. sapiens, N. gruberi*) and two species without (*A. thaliana, D. discoideum*) centrioles and motile cilia. Organisms known to have centrioles and motile cilia are indicated with centriole cartoons and include 4 clades of zoosporic ‘chytrid’ fungi (brown) that share a set of conserved centriole and cilium associated genes across functional categories (light green: centriole structure and duplication; green: pericentriolar material and centriole satellites; teal: centriole distal appendages; blue/purple: centriole accessory structures including subdistal appendage and basal feet as well as fibrous linkers).

Centrioles have a highly conserved, polarized structure that is intimately connected with their function. At the base of the centriole are structures associated with templating the centriole and tethering centrioles to other centrioles and organelles like the nucleus. At the distal tip of the centriole, the region where the axoneme extends from, are proteins that functionalize the structure to bind specific membranes and build the cilium. Chytrids have a large set of core conserved structural components as well as proteins involved in centriole duplication and elongation (e.g. CPAP, CEP120). They also have conserved genes for the distal appendage proteins that facilitate ciliogenesis and membrane docking, consistent with nine-fold symmetric densities in transmission electron micrographs (TEMs) that closely resemble those of animal centrioles (Longcore *et al*., 1999; Longcore and Simmons, 2020). Chytrids have some orthologs of other distal centriole components (Kumar and Reiter, 2021) (e.g. CC2D2A, TCHP, CEP19 and FOP) and fibrous linkers (e.g. TBCCD1, LRRCC1/VFL1, CCDC61,VFL3). Other electron-dense structures surrounding chytrid centrioles are similar within chytrid clades and have long been used for taxonomic classification but their function and molecular identity are not known as they do not closely resemble structures like the subdistal appendages of primary cilia or basal feet of motile cilia (Barr, 1981; Barr and Désaulniers, 1988; James *et al*., 2006; Kumar and Reiter, 2021).

In terms of microtubule structure and organization, chytrids have orthologs of the main tubulin family members including alpha, beta, gamma-tubulin as well as epsilon and zeta-tubulin (Chang *et al*., 2003; James *et al*., 2006; Turk *et al*., 2015; Kumar and Reiter, 2021). Most other fungi lack the expanded tubulin family members, zeta, epsilon, and delta-tubulin, while this ‘ZED’ module (Turk *et al*., 2015) is fully present in three of the four clades of zoosporic fungi, reduced in the Chytridiomycota, and absent in *Hyaloriphidium curvatum* that has lost cilia (Ustinova *et al*., 2000) (Figure S1). Chytrids have a conserved suite of adaptors involved in microtubule nucleation and organization including NEDD1, MZT1 and a larger family of gamma-tubulin complex proteins (GCPs) consistent with most other organisms, suggesting that the core microtubule nucleation module is conserved. Genes for pericentriolar material proteins are in general less conserved across eukaryotes than those of the centriole and similar to yeasts (e.g.

*S. cerevisiae* and *S. pombe*), chytrids have proteins with only short conserved domains of some animal PCM components (e.g. pericentrin, CDK5RAP2, CEP57 but not CEP192 or CEP152) involved in scaffolding pericentriolar material. Chytrids do not possess core modules of spindle pole bodies found in other fungi (e.g. SPC29, SPC42).

In sum, we have determined the genomic signature of chytrid centrioles and MTOCs. Chytrids have substantially more shared modules with centrioles than with spindle pole bodies and retain many conserved genomic signatures of centrosomes such as modules involved in organizing and scaffolding pericentriolar material and nucleating microtubules.

### Transcriptional signature of centriole maturation and ciliogenesis

Given that zoosporic fungi retain many of the genetic modules associated with centrosomes, we sought to characterize the transcriptional profile of MTOC-related genes and the structure of MTOCs over the lifecycle including mitosis and in the transition to ciliogenesis. To determine the transcriptional profiles associated with MTOC remodeling we isolated *R. globosum* at four timepoints after plating fresh zoospores (1.5 hour; germling, 13 hour; mitotic sporangia, 17.5 hour; ciliating sporangia, and 22 hour; new zoospores) and performed RNA-sequencing (Figure 3A, S2A). We found a core set of transcripts at all lifecycle stages, as well as hundreds of transcripts unique to these four lifecycle stages (Figure S1B). Overall we found the signature of an ordered transcriptional program underlying the different lifecycle stages (Rosenblum *et al*., 2012; Laundon *et al*., 2022) (Figure S2C-F).

**Figure 3.**
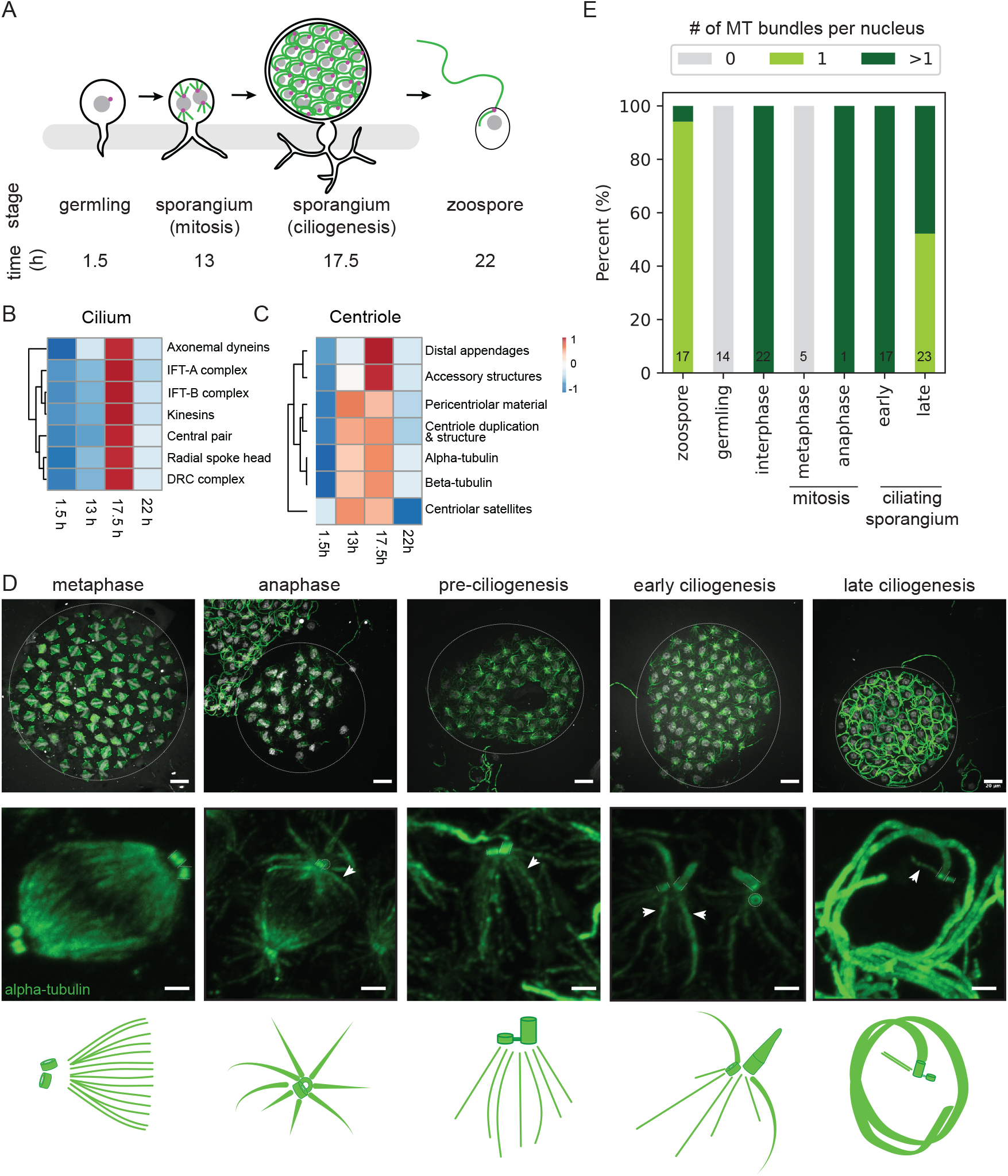
Centriole and MTOC architecture are remodeled during mitosis and ciliogenesis. A) Timepoints in the chytrid lifecycle used for RNA sequencing to probe key transitions (1.5 h early sporangium/germling, 13 h mitotic sporangium, 17.5 h ciliating sporangium, 22 h new zoospores). B,C) Heatmap of average RNA transcript abundance (FPKM, Fragments Per Kilobase of transcript per Million mapped reads) for groups of cilium and centriole related genes at each timepoint. Cilium related transcripts are upregulated at 17.5 hours during ciliogenesis. Centriole related transcripts are upregulated from 13 to 17.5 hours during mitosis and ciliogenesis while centriole accessory structures associated with distal maturation are specifically upregulated at ciliogenesis. D) Expansion microscopy characterization of chytrid MTOCs during mitosis and ciliogenesis (DAPI; white, alpha-tubulin; green). Scale bars 20 μm expanded (expansion factor ∼4.5 fold) and 2 μm for insets. Insets and summary cartoons show a single nucleus and associated cytoskeleton. Centrioles are short and orthogonal during mitotic divisions and oriented end on to the spindle pole. One centriole elongates at the start of ciliogenesis. MTOC architecture changes over the lifecycle from no astral microtubules at metaphase, large microtubule arrays in anaphase and during ciliogenesis, and a single microtubule bundle before zoospore release. E) Fraction of sporangia with each class of microtubule architecture at different lifecycle stages (n= number of sporangia from 3-5 independent experiments).

Ciliating sporangia have a strong transcriptional signature associated with ciliogenesis (Figure 3B) where components of the axoneme and ciliary beating are upregulated. We also looked for the signature of centrioles and their accessory structures, informed by our bioinformatic mapping (Figure 2, Figure 3C). Centriole-associated transcripts are upregulated during the mitotic cycles and ciliogenesis. Specifically, distal appendage components and subdistal appendage/basal foot components are strongly upregulated during ciliogenesis compared to mitosis, suggesting that, unlike in cycling mammalian cells, chytrid centrioles likely gain distal appendages just before ciliogenesis. This is consistent with the lack of distal appendage-like densities in TEM images of centrioles in mitotic chytrids (Longcore *et al*., 1999). We do not observe expression level differences in pericentriolar material components, which is consistent with post-translational regulation of PCM expansion common in other systems. Overall, we find ordered transcriptional programs associated with key lifecycle transitions into mitosis and ciliogenesis and specific modules associated with centriole remodeling and maturation.

### Microtubule organizing center remodeling from mitosis to ciliogenesis

In systems that form cilia, MTOCs often are remodeled, for example the centriole must change from being the center of a mitotic or interphase centrosome, to the base of the new cilium. Across eukaryotes, the centriole can continue to serve as an MTOC that patterns microtubules in the cytoplasm while also acting as the base of the cilium. Often in this arrangement, centrioles at the base of cilia are associated with ‘rootlet’ microtubule bundles that emanate from the side of specific microtubules in the centriole barrel (Azimzadeh, 2021). This is distinct from the type of MTOC structure that a centriole participates in as the center of a centrosome, where an aster of microtubules radiates from the PCM, rather than binding to specific regions of the centriole. In chytrids, microtubules have been observed radiating from centrioles at the base of cilia (James *et al*., 2006) as well as from centrioles at the spindle pole during mitosis, but the formation and structure of these MTOCs has not been systematically characterized. To determine the morphology of the mitotic MTOC in chytrids and how this structure changes at the end of the mitotic cycles to re-form cilia, we had to overcome the challenge of visualizing centriole structure using standard immunofluorescence due to the small size and low permeability of chytrid sporangia. Thus, we turned to ultrastructure expansion microscopy (U-ExM) (Gambarotto *et al*., 2019), a technique that physically enlarges a sample prior to antibody staining giving ∼4-5x increase in spatial resolution as well as improved permeabilization and lower background. This is the first use of expansion microscopy in zoosporic fungi and this technique is just beginning to be used in other fungal species (Götz *et al*., 2020; Hinterndorfer *et al*., 2022).

Using U-ExM, we mapped MTOC morphology during mitosis and ciliogenesis (Figure 3D). During mitosis, spindles form with one centriole in a pair oriented end-on to the spindle. This end-on arrangement has also been observed in other organisms, like chytrids, with semi-open mitoses (Powell, 1980; Shah *et al*., 2023). In late metaphase, centriole pairs lack bundles of astral microtubules which then reappear in early anaphase as spindle microtubules shorten (Figure 3D). The microtubules at the spindle pole resemble the radial array of microtubules found in animal centrosomes with symmetric distribution but no clear stereotyped orientation with respect to the centriole. At the end of the mitotic cycles, one centriole elongates to form the basal body (Figure 3D) (Lowry and Roberson, 1997; Lowry *et al*., 2004) and centrioles have a parallel orientation with bridging centrin signal (Figure 3D, Figure S3C-F). This remodeling of centriole length and orientation occurs prior to changes in the microtubule bundle architecture. Then the centriole at the base of the cilium is surrounded by microtubule bundles that resemble rootlets and terminate at the edge of the centriole (Figure 3D,E). By the end of ciliogenesis, there is a reduction in the complexity of the microtubule organization until only a single bundle of microtubules emanates from each basal body in the nascent zoospores prior to their release from the sporangium (Figure 3D).

Chytrid zoospore microtubules appear morphologically similar to microtubule rootlets of other ciliated eukaryotes (Barr, 1981; James *et al*., 2006; Yubuki and Leander, 2013; Azimzadeh, 2014). The transitions in the chytrid microtubule architecture that we observe during the lifecycle appear similar to that of the slime mold *Physarum polycepharum*, which in its amoeboid lifecycle alternates between a ciliated cell with centriole and rootlet microtubules and a mitotic cell type with a centrosome-like MTOC (Wright *et al*., 1988). The function of the microtubule bundles in chytrid zoospores and sporangia are not known. The microtubule bundles of the zoospore resemble rootlet microtubules in ciliated microeukaryotes like the green alga *Chlamydomonas reinhardtii* (Mittelmeier *et al*., 2011; Dutcher and O’Toole, 2016) and may play an analogous role (Galindo *et al*., 2021) in cell polarity and positioning of organelles or in ciliogenesis to scaffold intracellular or intraflagellar transport. Yet, unlike those of *C. reinhardtii*, chytrid microtubule organization is extensively remodeled to also adopt a geometry resembling a centrosomal aster at some stages of mitosis. In syncytial Drosophila embryos, centrosomal microtubule asters ensure proper nuclear spacing and organization of organelles (Lv *et al*., 2021), although the lack of microtubule asters during metaphase in chytrids suggests that these may not be required at all times to ensure proper spacing. In plants, microtubules often play a key role in cytokinesis, but depolymerization of the microtubule network in the post-mitotic sporangia of zoosporic fungus *Allomyces macrogynus* did not disrupt the analogous process of cellularization (Fisher *et al*., 2000).

### Centrioles shorten but are not degraded during ciliary disassembly

The cilium that is assembled at the end of mitosis enables the motility of free-swimming zoospores that are released from the sporangium. However, after chytrid zoospores attach to a substrate or host, they retract their cilium before progressing through their lifecycle (Koch, 1968; Berger *et al*., 2005; Medina and Buchler, 2020). During this transition the centriole must be liberated from the attached cilium to participate as an MTOC during mitosis. This problem is common to all cells that transition from a ciliated state to a non-ciliated state, from unicellular organisms to animal ciliated cells that participate in the cell cycle. Previously we characterized in detail the timeline of chytrid ciliary disassembly and found that over ∼2 hours cilia are retracted and axonemes are degraded (Venard *et al*., 2020). Using U-ExM, we were now able to characterize the structure and position of centrioles during ciliary retraction and degradation.

In expanded zoospores (Figure 4A), we observed centriole pairs with one long and one short centriole consistent with the architecture observed by EM (Barr and Désaulniers, 1988; Powell *et al*., 2019) (Figure S3A). In most chytrid species, zoospore centrioles have asymmetric length and a parallel orientation (Barr, 1981; Barr and Désaulniers, 1988; Venard *et al*., 2020). After retraction, we found that the cilium-centriole complex was coiled inside the cell body as the axoneme frayed (Figure 4B). Centrioles separated from the degrading axoneme and remained parallel, apparently linked by centrin-fibers (Figure 4C, S2). Instead of being degraded along with the axoneme, centrioles persisted and shortened, forming a pair of centrioles with equal length (Figure 4C,D). The centriole region that remained was enriched in centrin, consistent with the distribution in the proximal chytrid centriole barrel and with TEMs that show proximal structures like the cartwheel (Guichard *et al*., 2018) are present in both chytrid centrioles in a pair during mitosis (Barr, 1981; Barr and Désaulniers, 1988; Kalbfuss and Gönczy, 2023)(Figure S3A). Once the mitotic cycle began, centrioles of the coenocyte were short and orthogonal, and lacked bridging centrin fibers (Figure 4E, 3D, S3A-F). By analogy to the centriole cycle of animal cells, this may represent the engaged state (Nigg and Stearns, 2011) which is then followed by disengagement at the end of mitosis.

**Figure 4.**
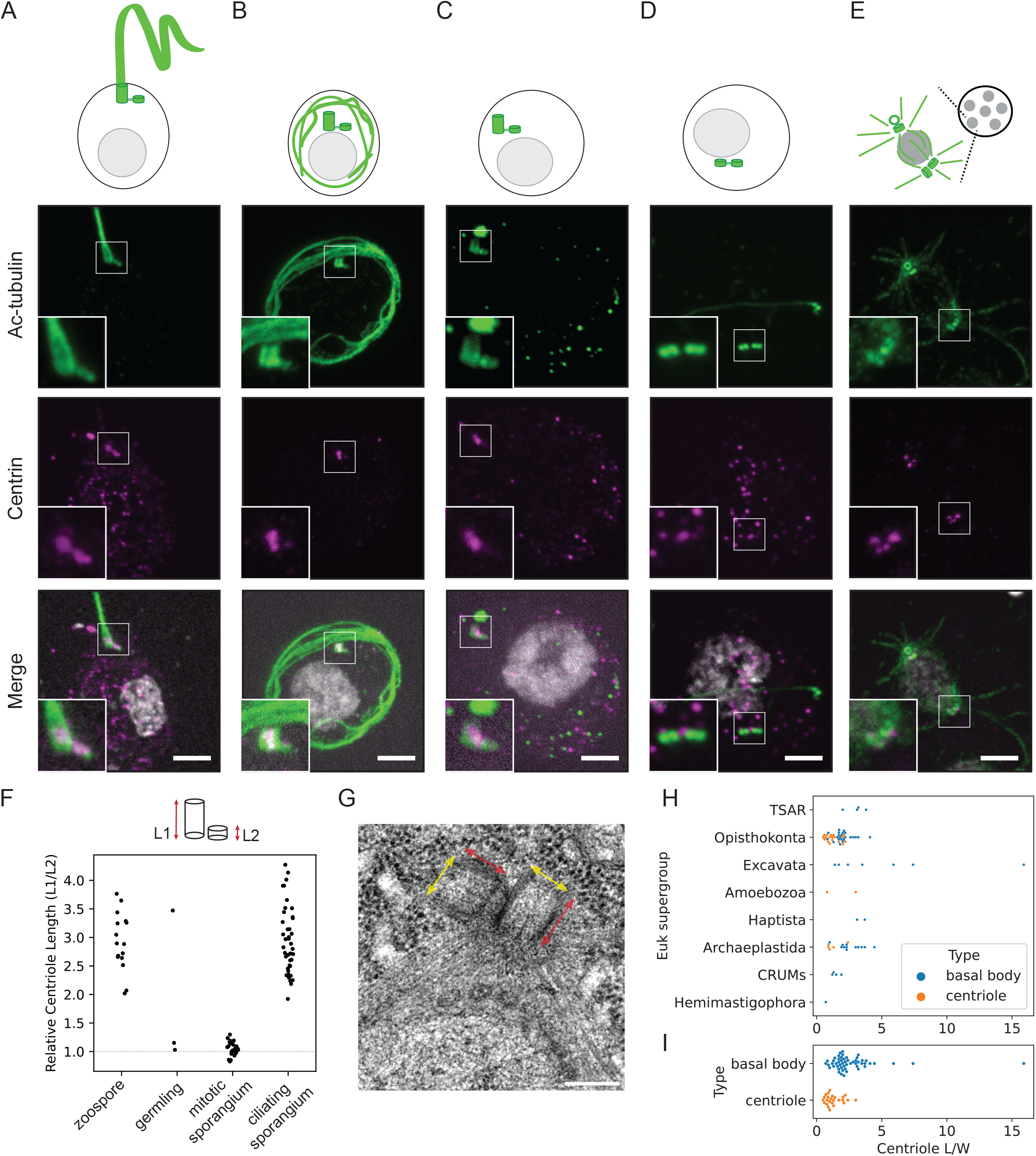
Chytrid fungi detach and shorten centrioles during axoneme degradation. A-E) Expansion microscopy characterization of chytrid centriole structure during the transition from basal body to mitotic MTOC (acetylated alpha-tubulin; green, centrin; magenta, DAPI/Hoechst; white). Scale bars 10 μm expanded (expansion factor ∼4.5 fold). Insets and summary cartoons show an individual centriole pair from the matched sporangium. Centrioles are long and parallel in the basal body and then after the centriole pair detaches from the axoneme, the longer centriole shortens in the germling while remaining parallel in orientation, linked by centrin fibers. During the mitotic cell cycles in the sporangium the centrioles are both short, equal in length, and orthogonal to each other. F) Quantifying the relative centriole length of each centriole pair over the lifecycle shows that in zoospores and the nascent zoospores of the ciliating sporangium, there is asymmetry in centriole length, while in the germling and mitotic sporangia centrioles are short and equal in length. n = 16, 3, 26, 41 sporangia respectively from 3-5 independent experiments. G) TEM image of mitotic *R. globosum* centriole pair showing equal length (red) and width (yellow) centrioles oriented orthogonally. H) Quantification of centriole length in basal body (blue) or centriole (orange) in a range of ciliated eukaryotic species (n=78 centrioles, 51 species). I) Mean centriole length of basal bodies (blue) versus centrioles (orange) pooled across eukaryotic species in (H) (n=78 centrioles, 51 species).

Reports of centriole shortening as we observed in chytrids are rare (Manton, 1964) as centrioles are normally quite stable structures (Kochanski and Borisy, 1990). In most contexts of cilium disassembly (Liang *et al*., 2016; Mirvis *et al*., 2018), cells separate the axoneme and centriole by resorbing the axoneme within an intact ciliary compartment or fully degrade both and synthesize new centrioles *de novo* (e.g in *Naegleria*) (Fulton and Dingle, 1971; Woglar *et al*., 2024). Centriole remodeling/reduction is most commonly seen during spermiogenesis (Avidor-Reiss and Fishman, 2019) which involves flaring and loss of the barrel structure, unlike the shortening of the barrel that we observed in chytrid centrioles. The mechanism that might protect a portion of the centriole during degradation of the axoneme in the cytoplasm is not known (Kalbfuss and Gönczy, 2023) but could result from specific post-translational modifications that protect the centriole or the stabilization of the proximal centriole through the cartwheel or inner centriole scaffold proteins (Le Guennec *et al*., 2020).

### Centriole structure and remodeling in evolution

The barrel structure of the centriole is highly conserved in evolution, yet the centriole shortening we observed during cilia disassembly in chytrids was striking. We wondered whether this phenomenon of a ‘short’ centriole with roughly equal length and width is an outlier or whether this is revealing some fundamental aspect of centriole biology that is conserved in eukaryotic evolution. We performed a meta-analysis of TEMs of diverse eukaryotic centrioles (Jana, 2021) across all major eukaryotic phyla (Figure 4G,H, Figure S4) and found that the average centriole aspect ratio (length/width) was 2.2 ± 0.2 (N=78, mean ± SEM). The ancestral centriole of the last eukaryotic common ancestor is thought to function exclusively as a basal body (Carvalho-Santos *et al*., 2011; Mitchell, 2017). We find that centrioles as basal bodies are on average longer than in contexts where they serve as MTOCs separate from the cilium (Figure 4I).

This length difference may be consistent with a loss of constraint on centriole size or a lack of requirement for the distal portion of the centriole when there is no cilium. For example, in the early divisions of the *Drosophila* or *C. elegans* embryo, centrioles remain short, with doublet or singlet MTs respectively and a persistent cartwheel structure, and only attain their mature elongated form later in differentiated cell types that will form cilia (Callaini *et al*., 1997; Pelletier *et al*., 2006). This is also true during mammalian differentiation, for example during the development of olfactory sensory neurons, short immature centrioles elongate and gain appendages only in the mature neuron once it is ready to ciliate (Ching *et al*., 2022). It is possible that the ancestral minimal centriole unit (Azimzadeh, 2021) for transmission through cell division is a small portion of the proximal centriole that lacks distal structures. It is not known whether the centriole is required for the function of the mitotic MTOC in chytrids, but restricting distal centriole assembly to ciliogenesis could serve as a mechanism to ensure separation of MTOC functions between centrosome and basal body at different lifecycle stages.

## Conclusion

In this study we develop the chytrid fungus *Rhizoclosmatium globosum* as a powerful system for studying MTOC remodeling as it undergoes major transitions in centriole and MTOC structure over its synchronous coenocytic lifecycle. Using immunofluorescence, expansion microscopy, and RNA sequencing we observe substantial remodeling of centrioles and MTOC architecture with different lengths and accessory structures associated with the centrosome-like MTOC versus the basal body as well as dynamic reorganizations of microtubule bundles. The specific remodeling of centriole structure over the lifecycle at the time of ciliogenesis instead of during the cell cycle is quite different from the typical centriole maturation cycle of cultured animal cells. Further, the shortening of the long centriole in the transition from basal body to mitotic MTOC is a surprising and rare form of centriole structural change. Looking forward, new genetic transformation methods in chytrids (Medina *et al*., 2020; Swafford *et al*., 2020; Kalinka *et al*., 2023; Prostak *et al*., 2023) will allow us to probe the regulatory mechanisms that drive the dramatic remodeling of chytrid centrioles and MTOCs and shed light on fundamental mechanisms for cytoskeletal remodeling and how they emerged in eukaryotic evolution.

## Acknowledgements

This work was made possible through the National Institutes of Health grants 1R35GM130286 to T.S, F32GM142181 to A.F.L and Stanford’s NHGRI training grant 5T32HG000044-23 to C.M.V. Many thanks to Joyce Longcore for the gift of *R. globosum* (JEL800) and for her tireless commitment to chytrid biology and generosity in sharing knowledge about chytrid diversity and evolution. We would like to thank Katie Ching for assistance with imaging and John Perrino and the Stanford Cell Science Imaging Facility for assistance in TEM image acquisition made possible through ARRA award NIH ORIP:1 S10 OD028536-01. J.E.S. and L.F-L are CIFAR Fellows in the Fungal Kingdom: Threats and Opportunities Program. We thank members of the Stearns and Feldman labs, Tim James, Nick Buchler, and the participants of the University of Michigan Biological Station Chytrid Workshop 2022 for helpful discussions. We thank the Guichard and Hamel labs for sharing their U-ExM Workshop protocol.

## Author Contributions

Conceptualization, A.F.L, K.V. and T.S.; investigation, A.F.L., K.V., C.M.V., A.J.M.S., J.E.S.; formal analysis-A.S., L.K. F-L., resources: L.K. F-L, J.L.F., T.S., writing– original draft, A.F.L., K.V.; review and editing, A.F.L, J.L.F, T.S.; visualization, A.F.L, A.S., K.V., supervision, T.S.

## Figure legends

**Figure S1: Centriole and centrosome gene conservation among chytrid species** Individual species level OrthoFinder analysis, supporting the phylum level trends in Figure 2, showing orthologs of genes associated with different centriole and centrosome features across two outgroups (top, gray) that have centrioles and motile cilia and all of the individual species included in our dataset within the zoosporic clades of fungi (NEOCAL, Neocallimastigomycetes; MON, Monoblepharidomycetes; CHYTR, Chytridiomycetes; BLAST, Blastocladiomycota). Each colored circle denotes that there was at least one ortholog assignment in the species. *R. globosum*, the organism in this study, is highlighted in brown.

**Figure S2: Ordered transcriptional programs underly key cytoskeletal transitions of the chytrid life cycle**. A) RNA sequencing of three biological replicates were performed for each of 4 timepoints. Heatmaps of the Pearson correlation coefficients for each sample show that biological replicates are highly correlated (scale 0, white to 1, black). B) Venn diagram showing unique and shared transcripts between four different timepoints in the lifecycle (1.5h, 13h, 17.5h, 22h). C-F) Heatmaps of individual genes showing transcript levels at four timepoints in the lifecycle. The groupings in C and D correspond to the individual genes averaged in Figure 3B,C. Groupings in E and F confirm that expected transcripts associated with DNA replication and channels and transporters are upregulated in timepoints with maturing sporangia and zoospore/germling respectively.

**Figure S3: Centriole size, polarity, and orientation during the chytrid life cycle**. A) TEM of orthogonal pair of centrioles from *R. globosum* (200 nm scale) showing presence of cartwheel (red arrow) in short centrioles (blue arrow denotes centriole length). B) Centriole width in micrometers measured from U-ExM gels (n=200 centrioles) versus TEM (n=11 centrioles) showing approximate 5 fold expansion factor C) Quantification of centriole pair orientation over the chytrid lifecycle (light blue: parallel, dark blue: orthogonal, gray: intermediate) showing that orthogonally oriented centrioles are only found in mitotic sporangia. D) Immunofluorescence images of expanded chytrid centriole pairs (matching insets in Figure 4A-D) showing alpha-tubulin (green) and centrin (magenta). E,F) Linescans of fluorescence intensity corresponding to images in (D) along width (blue) and length (yellow) of centriole pairs as diagrammed in each cartoon.

**Table S1: Centriole dimensions from electron micrographs**.

Meta-analysis of centriole length and width (n=78 centrioles) measured from TEM in published literature with corresponding species and cell type information (n=51 species), associated eukaryotic supergroup, unit of measure, and reference.

## Materials and Methods

### Culture and maintenance of chytrid strains

*Rhizoclosmatium globosum* strain JEL800 (*Rg*) were obtained from J. Longcore (University of Maine, Orono, ME). *Rg* cultures were grown on solid PmTG medium (0.1% peptonized milk, 0.1% tryptone, 0.5% glucose and 1% agar)(Donald J. S. Barr, 1986). For long term storage (6 months), *Rg* cultures were frozen and stored at −80 °C or under LN_2_(Boyle *et al*., 2003). Every 6 months, samples were freshly thawed and passaged twice onto solid medium for use in subsequent experiments.

Zoospores were isolated by adding PmTg liquid medium to 2 to 3-day old cultures for 3-5 min and then collecting the supernatant. The supernatant was checked under a light microscope for the absence of sporangia, and the concentration of zoospores was measured using a hemocytometer. The zoospores were then concentrated by spinning at 6000 RCF for 15s. The pelleted zoospores were resuspended, and 2×10^5^ spores were plated and incubated at 23°C for 1-24 hours depending on the experiment.

### Live cell imaging

Phase contrast images of the chytrid lifecycle (Figure 1) were acquired with a Keyence microscope with a Nikon 20x/ NA objective. Images were acquired for 3 z planes with 1 micron spacing every 5 minutes for 24 hours and the z-plane of best focus was chosen at each timepoint for subsequent analysis.

### Immunofluorescence

Samples were fixed by the addition of 1.5 mL of 4% formaldehyde directly to the plate. A cell scraper was used to gently dissociate cells from the plate. The fixed cells were transferred into an Eppendorf tube, incubated for 10 min at room temperature, and resuspended in PBS. In order to remove the cell wall, all time points other than 0 min were treated with 50 μg/mL chitinase (Sigma-Aldrich, Cat# C6137) in 20 mM potassium phosphate buffer pH 6.0 for 2h at 45°C. Cells were spun down onto poly-L-lysine coated coverslips, post-fixed 20 min in −20°C methanol, and blocked with PBS-BT (1X phosphate-buffered saline, 3% bovine serum albumin, 0.1% Triton-X 100, 0.02% sodium azide) for 1h. The cells were then stained for 2h at room temperature with different primary antibodies, and the concentrations used were: acetylated tubulin 6-11B-1 (Sigma-Aldrich, Cat# T6793;; RRID:AB_609894) 1:1000, α-tubulin (Sigma-Aldrich, Cat# T6199; RRID:AB_477583) 1:1000, centrin (EMD Millipore, Cat# 04-1624) 1:100. This was followed by incubation with relevant secondary antibodies with 5 μg/mL DAPI for 1 h before mounting into Mowiol on a slide for imaging. The different secondary antibodies (all were in a 1:1000 concentration in PBS-BT) used were: Alexa Fluor 488 Goat anti-mouse IgG2b antibody (Thermo Fisher Scientific, Cat# A-21141; RRID:AB_2535778) and Alexa Fluor 647 Goat anti-mouse IgG1 antibody (Thermo Fisher Scientific, Cat#A-21240;RRID:AB_2535809).

### Ultrastructure Expansion Microscopy

For expansion microscopy, chytrid sporangia were permeabilized through two different freeze-cracking methods rather than with chitinase digestion. In solution, chytrids fixed in 4% PFA in PBS in a microcentrifuge tube for 15 min at room temperature were flash-frozen in a dewar of liquid nitrogen, then thawed in a 37C or 42C water bath for 30-60s for between 1 and 10 cycles. Alternatively for freeze-cracking in a thin plane, a ∼12mm circle was drawn with a hydrophobic pen on a glass slide and the center was coated with fresh 1% poly-l-lysine solution for 30 min and rinsed 10 times with water and allowed to air dry for ∼10 minutes. Then 5-7 ul of chytrids fixed in 4% PFA in PBS for 15 min at room temperature were added to the center of the slide and sandwiched with a 22×50 mm #1.5 coverslip (Fisher Scientific Inc. 12544B) perpendicular to the slide. This slide sandwich was dipped into liquid nitrogen in a dewar for 10 seconds or until it stopped bubbling. Then the eraser end of a pencil was used to crack the coverslip off the slide. These slides were post-fixed in 100% cold methanol in coplin jars for 10 minutes then rehydrated in PBS

Fixed and permeabilized cells were incubated either 5 hours at 37°C or overnight at 4°C in 0.5 ml freshly prepared acrylamide/formaldehyde solution (AA/FA, 1.4% formaldehyde, 2% acrylamide in PBS) either in a 24 well dish or in a microcentrifuge tube. Gelation was allowed to proceed in monomer solution (19% sodium acrylate, 10% acrylamide, 0.1% bis-acrylamide, 0.5% ammonium persulfate-APS, 0.5% TEMED) on ice for 10 minutes followed by 1 hour at 37°C in a sealed humid chamber and then coverslips were discarded. Gels were boiled at 95°C in 2 ml denaturation buffer (200 mM SDS, 200 mM NaCl, 50 mM Tris pH 9) for 1.5 hours. Denaturation buffer was removed, gels were washed with multiple water rinses and allowed to expand in water at room temperature overnight. Small circles (approximately 5 mm in diameter of each expanded gel) were excised and incubated with primary antibodies diluted in PBS-BT buffer (3% BSA, 0.1%Triton X-100 in PBS) on a nutator at 4°C overnight. For experiments in Figures 3 and 4, primary antibodies were acetylated tubulin 6-11B-1 (Sigma-Aldrich, Cat# T6793; RRID:AB_609894) 1:1000, α-tubulin (Sigma-Aldrich, Cat# T6199; RRID:AB_477583) 1:1000, centrin (EMD Millipore, Cat# 04-1624; RRID:AB_10563501) 1:100. The next day, gels were washed three times with PBS-BT buffer and incubated with 5 μg/mL DAPI diluted in PBS-BT and secondary antibodies diluted 1:1000 were Alexa Fluor 488 Goat anti-mouse IgG2b antibody (Thermo Fisher Scientific, Cat# A-21141; RRID:AB_2535778), Alexa Fluor 568 Goat anti-mouse IgG1 antibody (Thermo Fisher Scientific, Cat# A-21124; RRID:AB_2535766), Alexa Fluor 647 Goat anti-mouse IgG2a antibody (Thermo Fisher Scientific, Cat#A-21241; RRID:AB_2535810). Gels were rocked on a nutator at 4°C overnight. Gels were washed once with 1X PBS and three times with water, and placed in a glass-bottom, freshly poly-L-lysine treated 35mm dish with #1.5 coverslip bottom for imaging (MatTek corp. P35G-1.5-20-C).

### Confocal microscopy

For images in Fig 1, slides were imaged using a Leica SP8 scanning confocal microscope with constant exposures during experiments, images were collected using the LAS X program and deconvolved with the HyVolution mode using the in-built Huygens deconvolution software from SVI. For imaging expansion microscopy gels in Fig 3 and 4, samples were imaged with an Zeiss Axio Observer microscope (Carl Zeiss) with a CSU-W1 confocal spinning-disk head (Yokogawa Electric Corporation, Tokyo, Japan), PlanApoChromat 63×/1.4 NA oil objective, and a PRIME: BSI backside illuminated CMOS camera run with SlideBook 6 software (3i, Denver, CO). Excitation lasers were 405nm, 488nm, 561nm, and 640nm. Z-stacks were acquired with 0.4 micron spacing.

### Transmission Electron Microscopy

For TEM, samples were fixed by addition of 1 ml of 2% glutaraldehyde and 4% paraformaldehyde in 0.1 M sodium cacodylate buffer, pH 7.2 on ice for 1 h. After fixation, the cells were spun down 2 min at 6k RPM and resuspended in warm 10% gelatin in PBS for 5 min at 37C. Samples were post fixed with 1ml of 1% osmium tetroxide for 1h on ice. The pellet was washed five times in water, followed by dehydration in an ethanol/water concentration series (50% to 100%). The samples were then washed in a series of propylene oxide/Epon concentrations followed by embedding in Epon. Ultrathin sections were stained with uranyl acetate and lead citrate and analyzed on a JEOL JEM1400 transmission electron microscope.

### RNA-sequencing

RNA-sequencing was done for *Rg* grown for 1.5 h, 13 h, 17.5 h, and 22 h after plating zoospores onto solid PmTG medium. We included three biological replicates for each time point. For each time point, 1 ml of 2×10^5^ zoospores/ml were plated on three 15 cm Petri dishes containing 1% PmTG solid medium and incubated at room temperature. Total RNA was extracted at the desired time points using Trizol reagent (Thermo Fisher Scientific, Cat# 15596026) following the manufacturer’s protocol. This was followed by DNAseI treatment (New England Biolabs, Cat# M0303S) and then re-extraction with Trizol. The quality of the total RNA sample was checked using a nanodrop; both A260/280 and A260/230 ratios were above 1.8.

The samples were sent to Novogene Corporation (https://en.novogene.com) for sequencing where they first analyzed RNA quality by Bioanalyzer (Agilent, USA) and then sequenced the samples using the NovaSeq 6000 (Illumina, San Diego, CA, USA), resulting in 150 bp paired-end sequences. Assembly of the sequencing data was performed by Novogene as follows: The raw data was filtered, and the sequences were aligned against the *Rg* reference genome using the program HISAT2. Gene expression levels were analyzed using HTseq software. Differentially expressed genes were identified with DESeq software. Volcano plot, heat map, GO enrichment, KEGG pathway analysis, and protein-protein interaction network analysis were also performed by Novogene.

For creating Venn diagrams (Fig S2B), we filtered for significantly expressed genes above 1 FPKM and input this list to the web tool: http://bioinformatics.psb.ugent.be/webtools/Venn/. Differential analysis of gene expression at each time point was done by manually analyzing the highly expressed genes at each time point for patterns using GO terms. For the grouped gene categories, ClustVis(Metsalu and Vilo, 2015) was used to generate heatmaps. The FPKM values were ln(x) transformed, rows were centered, and unit variance scaling was applied to rows. For creating the heatmap of the individual genes, no clustering was used (Figure S2), whereas rows and columns were clustered using Euclidean distance and average linkage for the averaged values of gene groups (Figure 3B,C).

### OrthoFinder Analysis

We used a combination of bioinformatic programs to identify the distribution of centriole and centrosome-related-protein orthologs across non-dikaryotic fungi. To do so, we downloaded the predicted genomes and proteomes of 213 non-dikaryotic fungi from JGI Mycocosm and 4 outgroups: Animals (*Homo sapiens*), Plants (*Arabidopsis thaliana*), *Naegleria*, and *Dictyostelium*. To guard against false positives, we used a number of quality control steps on the data, resulting in conservative estimates of orthology. First, we removed all datasets with BUSCO scores of less than 80% when compared to the default eukaryotic database (Manni *et al*., 2021). To generate an initial database of orthologs, we used Orthofinder (Emms and Kelly, 2019) with the remaining high-quality proteomes from JGI (accessed November 2023).

Orthofinder, however, significantly benefits from having a species tree imposed on the input data, but the existing phylogeny of non-dikaryotic fungi is still under frequent revision and does not include many of the species we have in this dataset. To account for this, we took an approach which allowed us to apply constraints from the existing literature and allow the data to inform unresolved relationships. We constructed a backbone phylogeny down to Order using the most currently reviewed phylogeny of non-dikaryotic fungi (Galindo *et al*., 2023) and then allowed the internal algorithms of Orthofinder to resolve phylogenetic relationships below the level of Order.

A limitation and strength of Orthofinder is that it only uses sequence data to determine orthology. While this makes results easier to interpret, it omits potentially important information that exists for many of these centriole and centrosome-related protein families. The hypothesized orthology of many centriole and centrosome-related proteins in more well studied organisms leverages additional structural, experimental, and theoretical data. To account for these additional data frequently available in Metazoa and Dikaryotic fungi but conspicuously absent in non-Dikarya, we manually assembled putatively-orthologous bait clusters with genes currently hypothesized to be orthologs based on data above the sequence level. Bait protein sequences were downloaded from UNIPROT (https://www.uniprot.org/) as FASTA. Baits include a list of common centriole and centrosome genes(Keller *et al*., 2005; Carvalho-Santos *et al*., 2010, 2011; Firat-Karalar *et al*., 2014; Galindo *et al*., 2021). Human bait sequences were included as the minimal reference with the exception of zeta-tubulin where we used the *Xenopus laevis* sequence as there is no copy in humans. In order to determine broad orthology groupings we chose to use bait clusters with sequences from multiple organisms that included the two fungi with the best studied microtubule organizing centers: *S. pombe* and *S. cerevisiae*. Thus, we included any known orthologs of bait proteins from the yeasts *S. pombe* and *S. cerevisiae*. Bait clusters also included manually curated orthologs from three zoosporic fungi, *R. globosum, B. dendrobatidis*, and *S. punctatus* where there were significant hits from pHMMER homology prediction and/or reciprocal blastp (NCBI) using the human protein sequence as bait.

We then used these putatively-orthologous bait clusters to merge all Orthofinder orthogroups represented by species within the putatively-orthologous bait clusters into a single, new merged orthogroup. As expected, many of these putatively-orthologous bait clusters mapped to a single Orthofinder orthogroup, but others included two or three Orthofinder orthogroups together. In these cases, we combined Orthofinder orthogroups into a merged orthogroup. Lastly, to remove spurious inclusions of protein fragments and off-target inclusions, we imposed a sequence length filter that removed any sequence shorter than 50% of the length of the mean bait sequence length used to create the new merged orthogroup. Each merged orthogroup was then scored against each species in our database, creating a binarized presence/absence database from our conservative estimate of orthologous genes.

### Data Analysis

Lifecycle stages in interphase and mitosis were determined through a combination of sporangium diameter and DNA organization (Fig 1,3,4) as *R. globosum* cell architecture and lifecycle timing is highly stereotyped.

Centriole length and width in expansion microscopy images or transmission electron micrographs was measured manually with a 1 pixel wide line in a single z plane of best focus using the linescan feature in Fiji version 2.0.0-rc-68/1.52g (Figure 4F, S3B,D-F). Only centrioles where the entire barrel lay along a z-plane were measured to minimize aberrations. For the alpha-tubulin and centrin intensity linescans in Figure S3, a 5 pixel wide line either 2 microns in length or 4 microns in width were drawn across the pair of centrioles in each axis respectively.

In the TEM meta-analysis dataset (Figure 4H,I,S4), centriole length is more challenging to score fairly as there is a wide diversity of distal ciliary and transition zone densities. We chose fiducials that avoided the transition zone and were present in as much of the dataset as possible. We measured manually using the linescan feature in Fiji from the proximal to distal end of the centriole and defined the distal endpoint as either the exposed end in a centrosome or the proximal end of any terminal plate structure if present. In the absence of any terminal plate, we measured to the base of the ciliary pocket membrane invagination if present. To maximize the number of images we could include in the dataset we report centriole size as an aspect ratio of length divided by width as many TEM images did not contain scale bars or single microtubules of known size to use as a fiducial. We chose at minimum two species per eukaryotic clade across all major kingdoms as well as deep branching eukaryotes of uncertain position (Torruella *et al*., 2024) (e.g. excavates) in lineages that retain centrioles (Nabais *et al*., 2020) informed by literature reviews of centriole length across diverse organisms (Hodges *et al*., 2012; Jana, 2021).

Manual linescan measurements described above were made with Fiji and then imported as CSV files into Python for subsequent analysis. Custom Python scripts written as Jupyter Notebooks were used to generate all plots and are available upon request.

## References

Avidor-Reiss, T, and Fishman, EL (2019). It takes two (centrioles) to tango. J Reprod Fertil 157, 1741–7899.

Azimzadeh, J (2014). Exploring the evolutionary history of centrosomes. Philos Trans R Soc Lond B Biol Sci 369: 20130453.

Azimzadeh, J (2021). Evolution of the centrosome, from the periphery to the center. Curr Opin Struct Biol 66, 96–103.

Barr, DJS (1981). The phylogenetic and taxonomic implications of flagellar rootlet morphology among zoosporic fungi. Biosystems 14, 359–370.

Barr, DJS (1986). Allochytridium expandens Rediscovered: Morphology, Physiology and Zoospore Ultrastructure. Mycologia 78, 439–448.

Barr, DJS, and Désaulniers, N (1988). Precise configuration of the chytrid zoospore. Botany 66, 869–876.

Berger, L, Hyatt, AD, Speare, R, and Longcore, JE (2005). Life cycle stages of the amphibian chytrid Batrachochytrium dendrobatidis. Dis Aquat Organ 68, 51–63.

Boyle, DG, Hyatt, AD, Daszak, P, Berger, L, Longcore, JE, Porter, D, Hengstberger, SG, and Olsen, V (2003). Cryo-archiving of Batrachochytrium dendrobatidis and other chytridiomycetes. Dis Aquat Organ 56, 59–64.

Callaini, G, Whitfield, WGF, and Riparbelli, MG (1997). Centriole and Centrosome Dynamics during the Embryonic Cell Cycles That Follow the Formation of the Cellular Blastoderm in Drosophila. Exp Cell Res 234, 183–190.

Carvalho-Santos, Z, Azimzadeh, J, Pereira-Leal, JB, and Bettencourt-Dias, M (2011). Evolution: Tracing the origins of centrioles, cilia, and flagella. J Cell Biol 194, 165–175.

Carvalho-Santos, Z, Machado, P, Branco, P, Tavares-Cadete, F, Rodrigues-Martins, A, Pereira-Leal, JB, and Bettencourt-Dias, M (2010). Stepwise evolution of the centriole-assembly pathway. J Cell Sci 123, 1414–1426.

Chang, P, Giddings, TH, Jr, Winey, M, and Stearns, T (2003). Epsilon-tubulin is required for centriole duplication and microtubule organization. Nat Cell Biol 5, 71–76.

Ching, K, Wang, JT, and Stearns, T (2022). Long-range migration of centrioles to the apical surface of the olfactory epithelium. eLife 11: e74399.

Dutcher, SK, and O’Toole, ET (2016). The basal bodies of Chlamydomonas reinhardtii. Cilia 5, 18.

Emms, DM, and Kelly, S (2015). OrthoFinder: solving fundamental biases in whole genome comparisons dramatically improves orthogroup inference accuracy. Genome Biol 16, 157.

Emms, DM, and Kelly, S (2019). OrthoFinder: phylogenetic orthology inference for comparative genomics. Genome Biol 20, 238.

Firat-Karalar, EN, Sante, J, Elliott, S, and Stearns, T (2014). Proteomic analysis of mammalian sperm cells identifies new components of the centrosome. J Cell Sci 127, 4128–4133.

Fisher, KE, Lowry, DS, and Roberson, RW (2000). Cytoplasmic cleavage in living zoosporangia of Allomyces macrogynus. J Microsc 198, 260–269.

Fulton, C, and Dingle, AD (1971). Basal bodies, but not centrioles, in Naegleria. J Cell Biol 51, 826–836.

Galindo, LJ, López-García, P, Torruella, G, Karpov, S, and Moreira, D (2021). Phylogenomics of a new fungal phylum reveals multiple waves of reductive evolution across Holomycota. Nat Commun 12, 4973.

Galindo, LJ, Torruella, G, López-García, P, Ciobanu, M, Gutiérrez-Preciado, A, Karpov, SA, and Moreira, D (2023). Phylogenomics Supports the Monophyly of Aphelids and Fungi and Identifies New Molecular Synapomorphies. Syst Biol 72, 505–515.

Gambarotto, D, Zwettler, FU, Le Guennec, M, Schmidt-Cernohorska, M, Fortun, D, Borgers, S, Heine, J, Schloetel, J-G, Reuss, M, Unser, M, et al. (2019). Imaging cellular ultrastructures using expansion microscopy (U-ExM). Nat Methods 16, 71–74.

Gleason Frank H., Scholz Bettina, Jephcott Thomas G., van Ogtrop Floris F., Henderson Linda, Lilje Osu, Kittelmann Sandra, and Macarthur Deborah J. (2017). Key Ecological Roles for Zoosporic True Fungi in Aquatic Habitats. Micro Spect 5, FUNK–0038-2016.

Götz, R, Panzer, S, Trinks, N, Eilts, J, Wagener, J, Turrà, D, Di Pietro, A, Sauer, M, and Terpitz, U (2020). Expansion Microscopy for Cell Biology Analysis in Fungi. Front Microbiol 11, 574.

Guichard, P, Hamel, V, and Gönczy, P (2018). The rise of the cartwheel: Seeding the centriole organelle. Bioessays 40, 1700241.

Hinterndorfer, K, Laporte, MH, Mikus, F, Tafur, L, Bourgoint, C, Prouteau, M, Dey, G, Loewith, R, Guichard, P, and Hamel, V (2022). Ultrastructure expansion microscopy reveals the cellular architecture of budding and fission yeast. J Cell Sci 135.

Hodges, ME, Scheumann, N, Wickstead, B, Langdale, JA, and Gull, K (2010). Reconstructing the evolutionary history of the centriole from protein components. J Cell Sci 123, 1407–1413.

Hodges, ME, Wickstead, B, Gull, K, and Langdale, JA (2012). The evolution of land plant cilia. New Phytol 195, 526–540.

Ito, D, and Bettencourt-Dias, M (2018). Centrosome Remodelling in Evolution. Cells 7, 71.

James, TY, Letcher, PM, Longcore, JE, Mozley-Standridge, SE, Porter, D, Powell, MJ, Griffith, GW, and Vilgalys, R (2006). A molecular phylogeny of the flagellated fungi (Chytridiomycota) and description of a new phylum (Blastocladiomycota). Mycologia 98, 860–871.

Jana, SC (2021). Centrosome structure and biogenesis: Variations on a theme? Semin Cell Dev Biol 110, 123–138.

Kalbfuss, N, and Gönczy, P (2023). Towards understanding centriole elimination. Open Biol 13, 230222.

Kalinka, E, Brody, SM, Swafford, AJM, Medina, EM, and Fritz-Laylin, LK (2024). Genetic transformation of the frog-killing chytrid fungus Batrachochytrium dendrobatidis. Proc Natl Acad Sci U S A 121, e2317928121.

Keller, LC, Romijn, EP, Zamora, I, Yates, JR, 3rd, and Marshall, WF (2005). Proteomic analysis of isolated Chlamydomonas centrioles reveals orthologs of ciliary-disease genes. Curr Biol 15, 1090–1098.

Kochanski, RS, and Borisy, GG (1990). Mode of centriole duplication and distribution. J Cell Biol 110, 1599–1605.

Koch, WJ (1968). Studies of the Motile Cells of Chytrids. V. Flagellar Retraction in Posteriorly Uniflagellate Fungi. Am J Bot 55, 841–859.

Kumar, D, and Reiter, J (2021). How the centriole builds its cilium: of mothers, daughters, and the acquisition of appendages. Curr Opin Struct Biol 66, 41–48.

Laundon, D, Chrismas, N, Bird, K, Thomas, S, Mock, T, and Cunliffe, M (2022). A cellular and molecular atlas reveals the basis of chytrid development. Elife 11, e73933.

Laundon, D, and Cunliffe, M (2021). A Call for a Better Understanding of Aquatic Chytrid Biology. Front Fungal Biol 2, 708813.

Le Guennec, M, Klena, N, Gambarotto, D, Laporte, MH, Tassin, A-M, van den Hoek, H, Erdmann, PS, Schaffer, M, Kovacik, L, Borgers, S, et al. (2020). A helical inner scaffold provides a structural basis for centriole cohesion. Sci Adv 6, eaaz4137.

Levy, YY, Lai, EY, Remillard, SP, Heintzelman, MB, and Fulton, C (1996). Centrin is a conserved protein that forms diverse associations with centrioles and MTOCs in Naegleria and other organisms. Cell Motil Cytoskeleton 33, 298–323.

Liang, Y, Meng, D, Zhu, B, and Pan, J (2016). Mechanism of ciliary disassembly. Cell Mol Life Sci 73, 1787–1802.

Longcore, JE, Pessier, AP, and Nichols, DK (1999). Batrachochytrium dendrobatidis gen. et sp. nov., a chytrid pathogenic to amphibians. Mycologia 91, 219–227.

Longcore, JE, and Simmons, DR (2020). Chytridiomycota. eLS, 1–9.

Lowry, DS, Fisher, KE, and Roberson, RW (2004). Functional necessity of the cytoskeleton during cleavage membrane development and zoosporogenesis in Allomyces macrogynus. Mycologia 96, 211–218.

Lowry, DS, and Roberson, RW (1997). Microtubule organization during zoosporogenesis in Allomyces macrogynus. Protoplasma 196, 45–54.

Lv, Z, de-Carvalho, J, Telley, IA, and Großhans, J (2021). Cytoskeletal mechanics and dynamics in the Drosophila syncytial embryo. J Cell Sci 134, jcs246496.

Manni, M, Berkeley, MR, Seppey, M, Simão, FA, and Zdobnov, EM (2021). BUSCO update: Novel and streamlined workflows along with broader and deeper phylogenetic coverage for scoring of eukaryotic, prokaryotic, and viral genomes. Mol Biol Evol 38, 4647–4654.

Manton, I (1964). Observations on the Fine Structure of the Zoospore and Young Germling of Stigeoclonium. J Exp Bot 15, 399–411.

Medina, EM, and Buchler, NE (2020). Chytrid fungi. Curr Biol 30, R516–R520.

Medina, EM, Robinson, KA, Bellingham-Johnstun, K, Ianiri, G, Laplante, C, Fritz-Laylin, LK, and Buchler, NE (2020). Genetic transformation of Spizellomyces punctatus, a resource for studying chytrid biology and evolutionary cell biology. Elife 9, e52741.

Metsalu, T, and Vilo, J (2015). ClustVis: a web tool for visualizing clustering of multivariate data using Principal Component Analysis and heatmap. Nucleic Acids Res 43, W566–W570.

Mirvis, M, Stearns, T, and James Nelson, W (2018). Cilium structure, assembly, and disassembly regulated by the cytoskeleton. Biochem J 475, 2329–2353.

Mitchell, DR (2017). Evolution of Cilia. Cold Spring Harb Perspect Biol 9, a028290.

Mittelmeier, TM, Boyd, JS, Lamb, MR, and Dieckmann, CL (2011). Asymmetric properties of the Chlamydomonas reinhardtii cytoskeleton direct rhodopsin photoreceptor localization. J Cell Biol 193, 741–753.

Mondo, SJ, Dannebaum, RO, Kuo, RC, Louie, KB, Bewick, AJ, LaButti, K, Haridas, S, Kuo, A, Salamov, A, Ahrendt, SR, et al. (2017). Widespread adenine N6-methylation of active genes in fungi. Nat Genet 49, 964–968.

Nabais, C, Peneda, C, and Bettencourt-Dias, M (2020). Evolution of centriole assembly. Curr Biol 30, R494–R502.

Nigg, EA, and Stearns, T (2011). The centrosome cycle: Centriole biogenesis, duplication and inherent asymmetries. Nat Cell Biol 13, 1154–1160.

Pelletier, L, O’Toole, E, Schwager, A, Hyman, AA, and Müller-Reichert, T (2006). Centriole assembly in Caenorhabditis elegans. Nature 444, 619–623.

Piperno, G, LeDizet, M, and Chang, XJ (1987). Microtubules containing acetylated alpha-tubulin in mammalian cells in culture. J Cell Biol 104, 289–302.

Powell, MJ (1980). Mitosis in the Aquatic Fungus Rhizophydium spherotheca (Chytridiales). Am J Bot 67, 839–853.

Powell, MJ (2017). Chytridiomycota. In: Handbook of the Protists: Second Edition, Springer International Publishing, 1523–1558.

Powell, MJ, Letcher, PM, Davis, WJ, Lefèvre, E, Brooks, M, and Longcore, JE (2019). Taxonomic summary of Rhizoclosmatium and description of four new Rhizoclosmatium species (Chytriomycetaceae, Chytridiales). Phytologia 101, 139–163.

Prostak, SM, Medina, EM, Kalinka, E, and Fritz-Laylin, LK (2023). A guide to Agrobacterium-mediated transformation of the chytrid fungus Spizellomyces punctatus. Access Microbiol 5, 000566.v3.

Prostak, SM, Robinson, KA, Titus, MA, and Fritz-Laylin, LK (2021). The actin networks of chytrid fungi reveal evolutionary loss of cytoskeletal complexity in the fungal kingdom. Curr Biol 31, 1192–1205.

Reiter, JF, and Leroux, MR (2017). Genes and molecular pathways underpinning ciliopathies. Nat Rev Mol Cell Biol 18, 533–547.

Rosenblum, EB, Poorten, TJ, Joneson, S, and Settles, M (2012). Substrate-specific gene expression in Batrachochytrium dendrobatidis, the chytrid pathogen of amphibians. PLoS One 7, e49924.

Rosenblum, EB, Stajich, JE, Maddox, N, and Eisen, MB (2008). Global gene expression profiles for life stages of the deadly amphibian pathogen Batrachochytrium dendrobatidis. Proc Natl Acad Sci U S A 105, 17034–17039.

Shah, H, Olivetta, M, Bhickta, C, Ronchi, P, Trupinić, M, Tromer, EC, Tolić, IM, Schwab, Y, Dudin, O, and Dey, G (2024). Life-cycle-coupled evolution of mitosis in close relatives of animals. Nature 630, 116–122.

Swafford, AJM, Hussey, SP, and Fritz-Laylin, LK (2020). High-efficiency electroporation of chytrid fungi. Sci Rep 10, 15145.

Taillon, BE, Adler, SA, Suhan, JP, and Jarvik, JW (1992). Mutational analysis of centrin: an EF-hand protein associated with three distinct contractile fibers in the basal body apparatus of Chlamydomonas. J Cell Biol 119, 1613–1624.

Torruella, G, Galindo, LJ, Moreira, D, and López-García, P (2024). Phylogenomics of neglected flagellated protists supports a revised eukaryotic tree of life. Curr Biol 35, 1–10.

Turk, E, Wills, AA, Kwon, T, Sedzinski, J, Wallingford, JB, and Stearns, T (2015). Zeta-Tubulin Is a Member of a Conserved Tubulin Module and Is a Component of the Centriolar Basal Foot in Multiciliated Cells. Curr Biol 25, 2177–2183.

Ustinova, I, Krienitz, L, and Huss, VAR (2000). Hyaloraphidium curvatum is not a Green Alga, but a Lower Fungus; Amoebidium parasiticum is not a Fungus, but a Member of the DRIPs. Protist 151, 253–262.

Venard, CM, Vasudevan, KK, and Stearns, T (2020). Cilium axoneme internalization and degradation in chytrid fungi. Cytoskeleton 77, 365–378.

Woglar, A, Busso, C, Garcia-Rodriguez, G, Douma, F, Croisier, M, Knott, G, and Gönczy, P (2024). Mechanisms of axoneme and centriole elimination in Naegleria gruberi. EMBO Rep 1–22.

Wright, M, Albertini, C, Planques, V, Salles, I, Ducommun, B, Gely, C, Akhavan-Niaki, H, Mir, L, Moisand, A, and Oustrin, ML (1988). Microtubule cytoskeleton and morphogenesis in the amoebae of the myxomycete Physarum polycephalum. Biol Cell 63, 239–248.

Yubuki, N, and Leander, BS (2013). Evolution of microtubule organizing centers across the tree of eukaryotes. Plant J 75, 230–244.

Zhang, Y, and He, CY (2012). Centrins in unicellular organisms: functional diversity and specialization. Protoplasma 249, 459–467.

