## Supplemental Figures S1-3 and Table S1 for "Dynamic remodeling of centrioles and the microtubule cytoskeleton in the lifecycle of chytrid fungi"

### Figure S1: Centriole and centrosome gene conservation among chytrid species

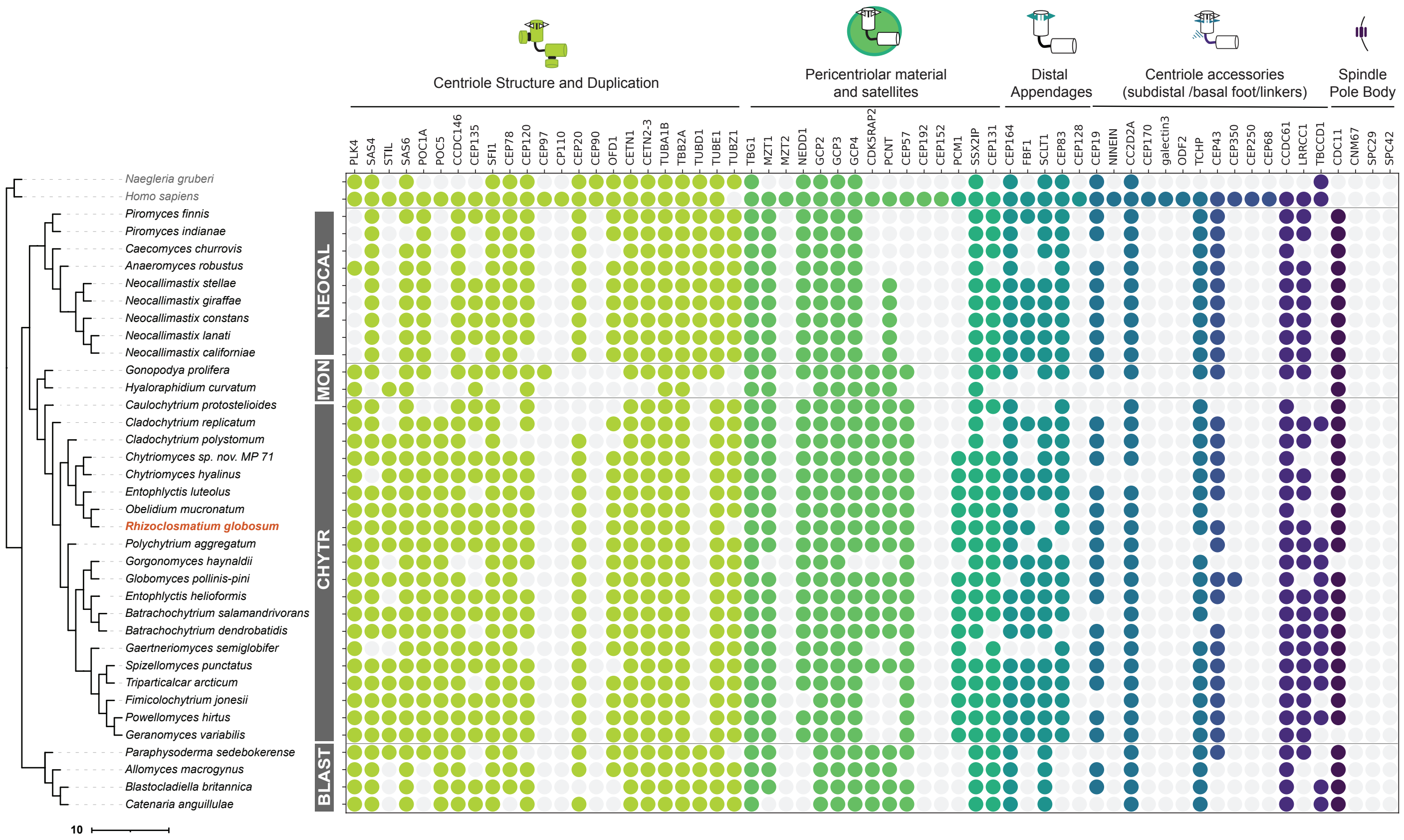

**Figure S2: Ordered transcriptional programs underly key cytoskeletal transitions of the chytrid life cycle**

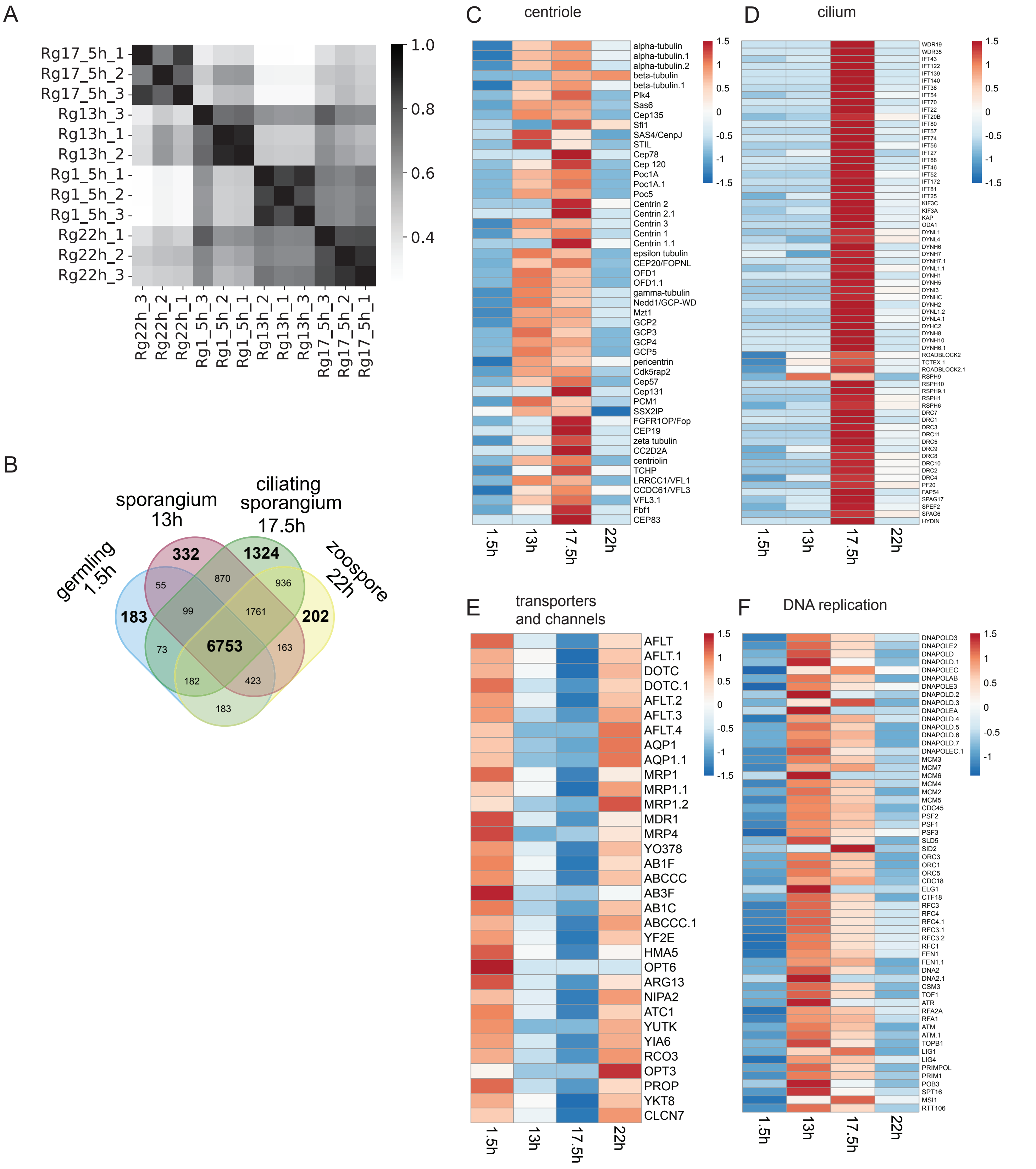

**Figure S3: Centriole size, polarity, and orientation change during the chytrid lifecycle**

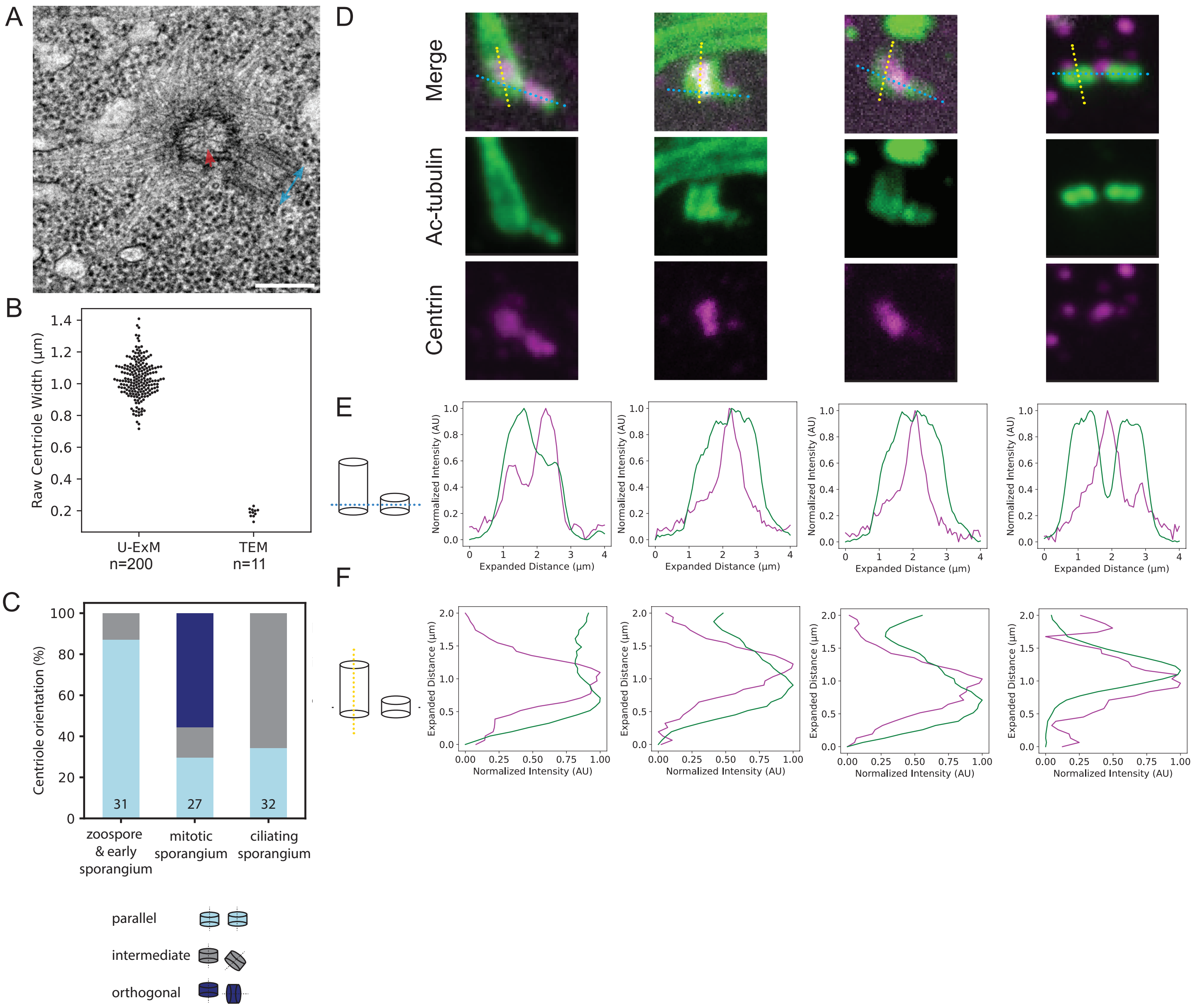

**Table S1: Centriole dimensions from electron micrographs.**

| NCBI ID | Organism | Cell Type | Supergroup | Type | Unit | Centriole length | Centriole width | Centriole L/W | Reference |
|---|---|---|---|---|---|---|---|---|---|
| 4785 | *Phytophthora cinnamomi* | zoospore | TSAR | basal body | px | 128 | 34 | 3.8 | (Hardham, 1987) |
| 4808 | *Blastocladiella emersonii* | sporangium | Opisthokonta | centriole | px | 41 | 57 | 0.7 | (Lessie and Lovett, 1968) |
| 4808 | *Blastocladiella emersonii* | sporangium | Opisthokonta | centriole | px | 72 | 60 | 1.2 | (Lessie and Lovett, 1968) |
| 4808 | *Blastocladiella emersonii* | zoospore | Opisthokonta | basal body | px | 234 | 79 | 3.0 | (Lessie and Lovett, 1968) |
| 5691 | *Trypanosoma brucei* | | Excavata | basal body | px | 167 | 68 | 2.4 | (Sherwin and Gull, 1989) |
| 5722 | *Trichomonas vaginalis* | | Excavata | basal body | nm | 616 | 174 | 3.5 | (Lee *et al.*, 2009) |
| 5741 | *Giardia intestinalis* | | Excavata | basal body | nm | 409 | 246 | 1.7 | (Nohynková *et al.*, 2006) |
| 5762 | *Naegleria gruberi* | | Excavata | basal body | nm | 281 | 198 | 1.4 | (Larson and Dingle, 1981) |
| 5791 | *Physarum polycephalum* | amoeba | Amoebozoa | centriole | px | 148 | 50 | 3.0 | (Gely and Wright, 1986) |
| 5791 | *Physarum polycephalum* | amoeba | Amoebozoa | centriole | px | 34 | 41 | 0.8 | (Gely and Wright, 1986) |
| 5888 | *Paramecium tetraurelia* | | TSAR | basal body | nm | 365 | 179 | 2.0 | (Bengueddach *et al.*, 2017) |
| 5911 | *Tetrahymena thermophila* | | TSAR | basal body | nm | 434 | 138 | 3.1 | (Bayless *et al.*, 2015) |
| 13221 | *Chrysotila carterae* | | Haptista | basal body | px | 137 | 45 | 3.1 | (Beech *et al.*, 1988) |
| 52552 | *Mallomonas splendens* | | TSAR | basal body | nm | 2278 | 701 | 3.2 | (Beech and Wetherbee, 1990) |
| 55998 | *Stigeoclonium sp.* | | Archaeplastida | centriole | nm | 177 | 199 | 0.9 | (Manton, 1964) |
| 55998 | *Stigeoclonium sp.* | | Archaeplastida | basal body | nm | 430 | 202 | 2.1 | (Manton, 1964) |
| 63628 | *Trichonympha agilis* | | Excavata | basal body | nm | 3112 | 196 | 15.9 | (Nazarov *et al.*, 2020) |
| 63628 | *Trichonympha agilis* | | Excavata | basal body | px | 294 | 40 | 7.4 | (Grimstone and Gibbons, 1966) |
| 64504 | *Allomyces arbusculus* | sporangium | Opisthokonta | centriole | px | 60 | 55 | 1.1 | (Renaud and Swift, 1964) |
| 64504 | *Allomyces arbusculus* | sporangium | Opisthokonta | basal body | px | 110 | 53 | 2.1 | (Renaud and Swift, 1964) |
| 65357 | *Albugo candida* | sporangium | Opisthokonta | centriole | nm | 183 | 220 | 0.8 | (Berlin and Bowen, 1964) |
| 65357 | *Albugo candida* | zoospore | Opisthokonta | basal body | nm | 620 | 240 | 2.6 | (Berlin and Bowen, 1964) |
| 70189 | *Olpidium brassicae* | zoospore | Opisthokonta | basal body | px | 135 | 83 | 1.6 | (Temmink and Campbell, 1969) |
| 81100 | *Breviata anathema* | | Opisthokonta | basal body | nm | 436 | 157 | 2.8 | (Heiss *et al.*, 2013a) |
| 81526 | *Monosiga ovata* | | Opisthokonta | basal body | nm | 376 | 183 | 2.1 | (Karpov, 2016) |
| 85706 | *Ancyromonas sp.* | | CRuMs | basal body | nm | 269 | 213 | 1.3 | (Heiss *et al.*, 2011) |
| 104085 | *Holomastigotoides sp.* | | Excavata | basal body | px | 562 | 95 | 5.9 | (Gibbons and Grimstone, 1960) |
| 109871 | *Batrachochytrium dendrobatidis* | zoospore | Opisthokonta | basal body | nm | 336 | 191 | 1.8 | (Longcore *et al.*, 1999) |
| 109876 | *Catenaria anguillulae* | sporangium | Opisthokonta | centriole | nm | 205 | 157 | 1.3 | (Ichida and Fuller, 1968) |
| 114254 | *Rhizidiomyces apophysatus* | sporangium | Opisthokonta | basal body | px | 36 | 11 | 3.2 | (Fuller and Reichle, 1965) |
| 190325 | *Collodictyon triciliatum* | | CRuMs | basal body | nm | 435 | 225 | 1.9 | (Brugerolle *et al.*, 2002) |
| 271399 | *Sorastrum sp.* | colony | Archaeplastida | centriole | px | 61 | 49 | 1.3 | (Marchant, 1974b) |
| 271399 | *Sorastrum sp.* | colony | Archaeplastida | basal body | px | 172 | 89 | 1.9 | (Marchant, 1974b) |
| 287562 | *Pediastrum boryanum* | zooid | Archaeplastida | centriole | px | 42 | 44 | 1.0 | (Marchant, 1974a) |
| 287562 | *Pediastrum boryanum* | zooid | Archaeplastida | basal body | px | 87 | 39 | 2.3 | (Marchant, 1974a) |
| 301696 | *Monoblepharis polymorpha* | zoospore | Opisthokonta | basal body | nm | 379 | 218 | 1.7 | (Mollicone and Longcore, 1994) |
| 329046 | *Rhizoclosmatium globosum* | zoospore | Opisthokonta | basal body | px | 93 | 55 | 1.7 | (Powell *et al.*, 2019) |
| 329046 | *Rhizoclosmatium globosum* | | Opisthokonta | centriole | px | 47 | 46 | 1.0 | This work. |
| 424538 | *Cricosphaera elongata* | | Haptista | basal body | nm | 1482 | 399 | 3.7 | (Henry *et al.*, 1991) |
| 529818 | *Thecamonas trahens* | | Opisthokonta | basal body | nm | 339 | 265 | 1.3 | (Heiss *et al.*, 2013b) |
| 906914 | *Chlamydomonas reinhardtii* | | Archaeplastida | basal body | nm | 358 | 183 | 2.0 | (Dutcher and O'Toole, 2016) |
| 2012329 | *Olpidium cucurbitacearum* | zoospore | Opisthokonta | basal body | nm | 328 | 177 | 1.9 | (Barr and Hadland-Hartmann, 1977) |

| NCBI ID | Organism | Cell Type | Supergroup | Type | Unit | Centriole length | Centriole width | Centriole L/W | Reference |
|---|---|---|---|---|---|---|---|---|---|
| 2012329 | *Olpidium cucurbitacearum* | zoospore | Opisthokonta | centriole | nm | 119 | 217 | 0.5 | (Barr and Hadland-Hartmann, 1977) |
| 2027450 | *Hemimastix amphikineta* | | Hemimastigophora | basal body | nm | 135 | 190 | 0.7 | (Foissner *et al.*, 1988) |
| 2823256 | *Caraotamonas croatica* | | CRuMs | basal body | nm | 184 | 126 | 1.5 | (Yubuki *et al.*, Nov-Dec 2023) |
| 2823262 | *Fabomonas mesopelagica* | | CRuMs | basal body | nm | 149 | 127 | 1.2 | (Yubuki *et al.*, Nov-Dec 2023) |
| 173374 | *Trentepohlia aurea* | | Archaeplastida | basal body | px | 131 | 57 | 2.3 | (Graham and McBride, 1975) |
| 13778 | *Nitella sp.* | | Archaeplastida | centriole | px | 53 | 59 | 0.9 | (Turner, 1968) |
| 13778 | *Nitella sp.* | | Archaeplastida | basal body | px | 152 | 47 | 3.2 | (Turner, 1968) |
| 3196 | *Marchantia sp.* | | Archaeplastida | basal body | px | 128 | 37 | 3.46 | (Carothers and Kreitner, 1968) |
| 67246 | *Aulacomnium palustre* | | Archaeplastida | centriole | nm | 448 | 190 | 2.4 | (Bernhard and Renzaglia, 1995) |
| 13804 | *Sphagnum sp.* | | Archaeplastida | basal body | px | 27.2 | 27.9 | 1.0 | (Manton, 1957) |
| 87755 | *Phaeoceros sp.* | | Archaeplastida | basal body | px | 50 | 21.6 | 2.3 | (Carothers *et al.*, 1977) |
| 3251 | *Lycopodium sp.* | | Archaeplastida | basal body | px | 111 | 25 | 4.44 | (Carothers *et al.*, 1975) |
| 3257 | *Equisetum sp.* | | Archaeplastida | basal body | px | 99 | 33 | 3 | (Duckett and Bell, 1977) |
| 3239 | *Psilotum sp.* | | Archaeplastida | basal body | nm | 684 | 186 | 3.7 | (Renzaglia *et al.*, 2001) |
| 9606 | *Homo sapiens* | breast epithelial | Opisthokonta | centriole | nm | 397 | 190 | 2.1 | (Lingle *et al.*, 1998) |
| 9606 | *Homo sapiens* | U2OS | Opisthokonta | centriole | nm | 382 | 189 | 2.02 | (Arquint *et al.*, 2012) |
| 9606 | *Homo sapiens* | RPE-1 | Opisthokonta | centriole | nm | 434 | 201 | 2.2 | (Marteil *et al.*, 2018) |
| 9606 | *Homo sapiens* | RPE-1 | Opisthokonta | basal body | nm | 342 | 158 | 2.2 | (Molla-Herman *et al.*, 2010) |
| 10090 | *Mus musculus* | olfactory epithelial | Opisthokonta | centriole | nm | 190 | 190 | 1 | (Falk *et al.*, 2015) |
| 10090 | *Mus musculus* | olfactory epithelial | Opisthokonta | basal body | nm | 412 | 190 | 2.2 | (Falk *et al.*, 2015) |
| 10090 | *Mus musculus* | photoreceptor | Opisthokonta | basal body | nm | 410 | 209 | 2.0 | (Falk *et al.*, 2015) |
| 10090 | *Mus musculus* | trachea | Opisthokonta | basal body | nm | 423 | 235 | 1.8 | (Burke *et al.*, 2014) |
| 7955 | *Danio rerio* | larval brain | Opisthokonta | basal body | nm | 259 | 154 | 1.7 | (Wilkinson *et al.*, 2009) |
| 7955 | *Danio rerio* | neuroepithelium | Opisthokonta | centriole | nm | 273 | 163 | 1.7 | (Dzafic *et al.*, 2015) |
| 9031 | *Gallus domesticus* | choroid plexus | Opisthokonta | basal body | nm | 1506 | 793 | 1.9 | (Stephen *et al.*, 2015) |
| 9031 | *Gallus domesticus* | choroid plexus | Opisthokonta | centriole | nm | 537 | 274 | 2.0 | (Stephen *et al.*, 2015) |
| 6239 | *Caenorhabditis elegans* | embryo | Opisthokonta | centriole | nm | 208 | 177 | 1.2 | (Pelletier *et al.*, 2006) |
| 7227 | *Drosophila melanogaster* | auditory neurons | Opisthokonta | basal body | nm | 196 | 86 | 2 | (Jana *et al.*, 2018) |
| 7227 | *Drosophila melanogaster* | spermatocyte | Opisthokonta | basal body | nm | 1667 | 410 | 4.1 | (Jana *et al.*, 2018) |
| 7227 | *Drosophila melanogaster* | larval wing disc | Opisthokonta | centriole | nm | 124 | 143 | 0.87 | (Franz *et al.*, 2013) |
| 7227 | *Drosophila melanogaster* | larval brain | Opisthokonta | centriole | nm | 93 | 157 | 0.59 | (Franz *et al.*, 2013) |
| 7227 | *Drosophila melanogaster* | olfactory neuron | Opisthokonta | basal body | nm | 173 | 208 | 0.83 | (Gottardo *et al.*, 2015) |
| 7227 | *Drosophila melanogaster* | gonioblast | Opisthokonta | basal body | nm | 168 | 187 | 0.9 | (Gottardo *et al.*, 2015) |
| 7227 | *Drosophila melanogaster* | spermatogone | Opisthokonta | basal body | nm | 341 | 198 | 1.72 | (Gottardo *et al.*, 2015) |
| 76800 | *Anisopteromalus calandrae* | larval cell | Opisthokonta | centriole | nm | 137 | 217 | 0.6 | (Uzbekov *et al.*, 2018) |
| 76800 | *Anisopteromalus calandrae* | spermatid | Opisthokonta | centriole | nm | 275 | 197 | 1.40 | (Uzbekov *et al.*, 2018) |

#### Supplemental References for Table S1


Arquint, C, Sonnen, KF, Stierhof, Y-D, and Nigg, EA (2012). Cell-cycle-regulated expression of STIL controls centriole number in human cells. J Cell Sci 125, 1342–1352.

Barr, DJS, and Hadland-Hartmann, VE (1977). Zoospore ultrastructure of *Olpidium cucurbitacearum* (Chytridiales). Can J Bot 55, 3063–3074.

Bayless, BA, Galati, DF, and Pearson, CG (2015). *Tetrahymena* basal bodies. Cilia 5, 1.

Beech, PL, and Wetherbee, R (1990). The Flagellar Apparatus of *Mallomonas splendens* (Synurophyceae) at Interphase and its Development During the Cell Cycle. J Phycol 26, 95–111.

Beech, PL, Wetherbee, R, and Pickett-Heaps, JD (1988). Transformation of the flagella and associated flagellar components during cell division in the coccolithophorid *Pleurochrysis carterae*. Protoplasma 145, 37–46.

Bengueddach, H, Lemullois, M, Aubusson-Fleury, A, and Koll, F (2017). Basal body positioning and anchoring in the multiciliated cell *Paramecium tetraurelia*: roles of OFD1 and VFL3. Cilia 6, 6.

Berlin, JD, and Bowen, CC (1964). Centrioles in the fungus *Albugo candida*. Am J Bot 51, 650–652.

Bernhard, DL, and Renzaglia, KS (1995). Spermiogenesis in the Moss *Aulacomnium palustre*. Bryologist 98, 52.

Brugerolle, G, Bricheux, G, Philippe, H, and Coffe, G (2002). *Collodictyon triciliatum* and *Diphylleia rotans* (*Aulacomonas submarina*) form a new family of flagellates (Collodictyonidae) with tubular mitochondrial cristae that is phylogenetically distant from other flagellate groups. Protist 153, 59–70.

Burke, MC, Li, F-Q, Cyge, B, Arashiro, T, Brechbuhl, HM, Chen, X, Siller, SS, Weiss, MA, O'Connell, CB, Love, D, *et al.* (2014). Chibby promotes ciliary vesicle formation and basal body docking during airway cell differentiation. J Cell Biol 207, 123–137.

Carothers, ZB, and Kreitner, GL (1968). Studies of spermatogenesis in the Hepaticae. II. Blepharoplast structure in the spermatid of *Marchantia*. J Cell Biol 36, 603–616.

Carothers, ZB, Moser, JW, and Duckett, JG (1977). Ultrastructural studies of spermatogenesis in the anthocerotales. Ii. The blepharoplast and anterior mitochondrion in *Phaeoceros laevis*: Later development. Am J Bot 64, 1107–1116.

Carothers, ZB, Robbins, RR, and Haas, DL (1975). Some ultrastructural aspects of spermatogenesis in *Lycopodium complanatum*. Protoplasma 86, 339–350.

Duckett, JG, and Bell, PR (1977). An ultrastructural study of the mature spermatozoid of*Equisetum*. Philos Trans R Soc Lond 277, 131–158.

Dutcher, SK, and O'Toole, ET (2016). The basal bodies of *Chlamydomonas reinhardtii*. Cilia 5, 18.

Dzafic, E, Strzyz, PJ, Wilsch-Bräuninger, M, and Norden, C (2015). Centriole amplification in zebrafish affects proliferation and survival but not differentiation of neural progenitor cells. Cell Rep 13, 168–182.

Falk, N, Lösl, M, Schröder, N, and Gießl, A (2015). Specialized cilia in mammalian sensory systems. Cells 4, 500–519.

Foissner, W, Blatterer, H, and Foissner, I (1988). The hemimastigophora (*Hemimastix amphikineta* nov. gen., nov. spec.), a new protistan phylum from gondwanian soils. Eur J Protistol 23, 361–383.

Franz, A, Roque, H, Saurya, S, Dobbelaere, J, and Raff, JW (2013). CP110 exhibits novel regulatory activities during centriole assembly in *Drosophila*. J Cell Biol 203, 785–799.

Fuller, MS, and Reichle, R (1965). The Zoospore and Early Development of *Rhizidiomyces apophysatus*. Mycologia 57, 946–961.

Gely, C, and Wright, M (1986). The centriole cycle in the amoebae of the myxomycete *Physarum polycephalum*. Protoplasma 132, 23–31.

Gibbons, IR, and Grimstone, AV (1960). On flagellar structure in certain flagellates. J Biophys Biochem Cytol 7, 697–716.

Gottardo, M, Callaini, G, and Riparbelli, MG (2015). The *Drosophila* centriole - conversion of doublets into triplets within the stem cell niche. J Cell Sci 128, 2437–2442.

Graham, LE, and McBride, GE (1975). The Ultrastructure of Multilayered Structures Associated with Flagellar Bases in Motile Cells of *Trentepohlia aurea* . J Phycol 11, 86–96.

Grimstone, AV, and Gibbons, IR (1966). The fine structure of the centriolar apparatus and associated structures in the complex flagellates *Trichonympha* and *Pseudotrichonympha*. Philos Trans R Soc Lond 250, 215–242.

Hardham, AR (1987). Microtubules and the flagellar apparatus in zoospores and cysts of the fungusPhytophthora cinnamomi. Protoplasma 137, 109–124.

Heiss, AA, Walker, G, and Simpson, AGB (2011). The ultrastructure of Ancyromonas, a eukaryote without supergroup affinities. Protist 162, 373–393.

Heiss, AA, Walker, G, and Simpson, AGB (2013a). The flagellar apparatus of *Breviata anathema*, a eukaryote without a clear supergroup affinity. Eur J Protistol 49, 354–372.

Heiss, AA, Walker, G, and Simpson, AGB (2013b). The microtubular cytoskeleton of the apusomonad *Thecamonas*, a sister lineage to the opisthokonts. Protist 164, 598–621.

Henry, M, Karez, CS, Rom, M, Gnassia-Barelli, M, and Puiseux-Dao, S (1991). Ultrastructural study and calcium and cadmium localization in the marine coccolithophorid *Cricosphaera elongata*. Mar Biol 111, 167–173.

Ichida, AA, and Fuller, MS (1968). Ultrastructure of mitosis in the aquatic fungus *Catenaria anguillulae*. Mycologia 60, 141–155.

Jana, SC, Mendonça, S, Machado, P, Werner, S, Rocha, J, Pereira, A, Maiato, H, and Bettencourt-Dias, M (2018). Differential regulation of transition zone and centriole proteins contributes to ciliary base diversity. Nat Cell Biol 20, 928–941.

Karpov, SA (2016). Flagellar apparatus structure of choanoflagellates. Cilia 5, 11.

Larson, DE, and Dingle, AD (1981). Isolation, ultrastructure, and protein composition of the flagellar rootlet of *Naegleria gruberi*. J Cell Biol 89, 424–432.

Lee, KE, Kim, JH, Jung, MK, Arii, T, Ryu, J-S, and Han, SS (2009). Three-dimensional structure of the cytoskeleton in *Trichomonas vaginalis* revealed new features. J Electron Microsc 58, 305–313.

Lessie, PE, and Lovett, JS (1968). Ultrastructural changes during sporangium formation and zoospore differentiation in *Blastocladiella emersonii*. Am J Bot 55, 220–236.

Lingle, WL, Lutz, WH, Ingle, JN, Maihle, NJ, and Salisbury, JL (1998). Centrosome hypertrophy in human breast tumors: implications for genomic stability and cell polarity. Proc Natl Acad Sci U S A 95, 2950–2955.

Longcore, JE, Pessier, AP, and Nichols, DK (1999). *Batrachochytrium dendrobatidis* gen. et sp. nov., a chytrid pathogenic to amphibians. Mycologia 91, 219–227.

Manton, I (1957). Observations with the electron microscope on the cell structure of the antheridium and spermatozoid of Sphagnum. J Exp Bot 8, 382–400.

Manton, I (1964). Observations on the Fine Structure of the Zoospore and Young Germling of *Stigeoclonium*. J Exp Bot 15, 399–411.

Marchant, HJ (1974a). Mitosis, Cytokinesis and Colony Formation in *Pediastrum boryanum*. Ann Bot 38, 883–888.

Marchant, HJ (1974b). Mitosis, Cytokinesis, and Colony Formation in the Green Alga *Sorastrum*. J Phycol 10, 107–120.

Marteil, G, Guerrero, A, Vieira, AF, de Almeida, BP, Machado, P, Mendonça, S, Mesquita, M, Villarreal, B, Fonseca, I, Francia, ME, *et al.* (2018). Over-elongation of centrioles in cancer promotes centriole amplification and chromosome missegregation. Nat Commun 9, 1258.

Molla-Herman, A, Ghossoub, R, Blisnick, T, Meunier, A, Serres, C, Silbermann, F, Emmerson, C, Romeo, K, Bourdoncle, P, Schmitt, A, *et al.* (2010). The ciliary pocket: an endocytic membrane domain at the base of primary and motile cilia. J Cell Sci 123, 1785–1795.

Mollicone, MRN, and Longcore, JE (1994). Zoospore ultrastructure of *Monoblepharis polymorpha*. Mycologia 86, 615–625.

Nazarov, S, Bezler, A, Hatzopoulos, GN, Nemčíková Villímová, V, Demurtas, D, Le Guennec, M, Guichard, P, and Gönczy, P (2020). Novel features of centriole polarity and cartwheel stacking revealed by cryo-tomography. EMBO J 39, e106249.

Nohynková, E, Tumová, P, and Kulda, J (2006). Cell division of *Giardia intestinalis*: flagellar developmental cycle involves transformation and exchange of flagella between mastigonts of a diplomonad cell. Eukaryot Cell 5, 753–761.

Pelletier, L, O'Toole, E, Schwager, A, Hyman, AA, and Müller-Reichert, T (2006). Centriole assembly in *Caenorhabditis elegans*. Nature 444, 619–623.

Powell, MJ, Letcher, PM, Davis, WJ, Lefèvre, E, Brooks, M, and Longcore, JE (2019). Taxonomic summary of *Rhizoclosmatium* and description of four new *Rhizoclosmatium* species (Chytriomycetaceae, Chytridiales). Phytologia 101, 139–163.

Renaud, FL, and Swift, H (1964). The Development of Basal Bodies and Flagella in *Allomyces arbusculus*. J Cell Biol 23, 339–354.

Renzaglia, KS, Johnson, TH, Gates, HD, and Whittier, DP (2001). Architecture of the sperm cell of *Psilotum*. Am J Bot 88, 1151–1163.

Sherwin, T, and Gull, K (1989). The cell division cycle of *Trypanosoma brucei brucei* : timing of event markers and cytoskeletal modulations. Philos Trans R Soc Lond B Biol Sci 323, 573–588.

Stephen, LA, Tawamie, H, Davis, GM, Tebbe, L, Nürnberg, P, Nürnberg, G, Thiele, H, Thoenes, M, Boltshauser, E, Uebe, S, *et al.* (2015). TALPID3 controls centrosome and cell polarity and the human ortholog KIAA0586 is mutated in Joubert syndrome (JBTS23). Elife 4, e08077.

Temmink, JHM, and Campbell, RN (1969). The ultrastructuse of *Olpidium brassicae*. TI. Zoospores. Canadian Journal of Botany, 227–231.

Turner, FR (1968). An ultrastructural study of plant spermatogenesis. Spermatogenesis in *Nitella*. J Cell Biol 37, 370–393.

Uzbekov, R, Garanina, A, and Bressac, C (2018). Centrioles without microtubules: a new morphological type of centriole. Biol Open 7.

Wilkinson, CJ, Carl, M, and Harris, WA (2009). Cep70 and Cep131 contribute to ciliogenesis in zebrafish embryos. BMC Cell Biol 10, 17.

Yubuki, N, Torruella, G, Galindo, LJ, Heiss, AA, Ciobanu, MC, Shiratori, T, Ishida, K-I, Blaz, J, Kim, E, Moreira, D, *et al.* (Nov-Dec 2023). Molecular and morphological characterization of four new ancyromonad genera and proposal for an updated taxonomy of the Ancyromonadida. J Eukaryot Microbiol 70, e12997.
